# RNA-targeting CRISPR-Cas13 Provides Broad-spectrum Phage Immunity

**DOI:** 10.1101/2022.03.25.485874

**Authors:** Benjamin A. Adler, Tomas Hessler, Brady F Cress, Vivek K. Mutalik, Rodolphe Barrangou, Jillian Banfield, Jennifer A Doudna

## Abstract

CRISPR-Cas13 proteins are RNA-guided RNA nucleases that defend against invasive phages through general, non-specific RNA degradation upon complementary target transcript binding. Despite being RNA nucleases, Cas13 effectors are capable of inhibiting the infection of dsDNA phages but have only been investigated across a relatively small sampling of phage diversity. Here, we employ a systematic, phage-centric approach to determine the anti-phage capacity of Cas13 and find LbuCas13a to be a remarkably potent phage inhibitor. LbuCas13a confers robust, consistent antiviral activity regardless of gene essentiality, gene expression timing or target sequence location. Furthermore, after challenging LbuCas13a with eight diverse *E. coli* phages distributed across *E. coli* phage phylogenetic groups, we find no apparent phage-encoded limits to its potent antiviral activity. In contrast to other Class 2 CRISPR-Cas proteins, these results suggest that DNA phages are generally vulnerable to Cas13a targeting. Leveraging this effective anti-phage activity, LbuCas13a can be used seamlessly as a counter-selection agent for broad-spectrum phage editing. Using a two-step phage editing and enrichment approach, we show that LbuCas13a enables markerless genome edits in phages with exceptionally high efficiency and precision, including edits as small as a single codon. By taking advantage of the broad vulnerability of RNA during viral infection, Cas13a enables a generalizable strategy for editing the most abundant and diverse biological entities on Earth.

## Introduction

CRISPR-Cas systems confer diverse RNA-guided antiviral and anti-plasmid adaptive immunity in prokaryotes^1^. CRISPR genomic loci record phage infections over time in the form of sequence arrays comprising foreign DNA sequences (spacers) flanked by direct repeats (DRs). Array transcription and processing generates CRISPR RNAs (crRNAs) that associate with one or more cognate Cas proteins to form ribonucleoprotein complexes capable of recognizing crRNA-complementary DNA or RNA^2^. Upon target binding, Cas effectors disrupt phage infection using DNA cleavage^3–5^, RNA cleavage^6^, secondary messenger production^7, 8^, or transcriptional silencing^9^. These programmable biochemical activities have had tremendous success as genome editing tools in bacteria and eukaryotes^10^.

Due to the coevolutionary arms race between phages and their target bacteria, phages encode direct and indirect inhibitors of CRISPR-Cas systems^11–14^, employ DNA compartmentalizing or masking strategies^15–19^, and manipulate DNA repair systems^20, 21^. In addition, phages use population-level strategies to overwhelm^22, 23^ and even destroy native CRISPR pathways^24^. This suite of active and passive DNA defense mechanisms has rendered it difficult to generalize the use of any single DNA-targeting CRISPR effector as a sequence-guided phage genome editing tool^25–28^.

Cas13 (formerly C2c2) effectors are RNA-guided RNA nucleases whose catalytic activity resides in two higher eukaryotic and prokaryotic nucleotide binding (HEPN) domains^6, 29^. Four Cas13 subtypes (a-d) differ by primary sequence and size as well as auxiliary gene association and extent of cis-versus trans-RNA cleavage activity^2^. Distinct from other single effector CRISPR-Cas systems, Cas13 is capable of conferring both individual- and population-level defense against phage infection^30^. Upon target RNA binding, Cas13 unleashes general, non-specific RNA degradation that arrests growth of the virocell to block infection progression, thereby limiting infection of neighboring cells^30^. Since all known viruses produce RNA^31^, Cas13 is capable of inhibiting dsDNA phages, primarily shown through studies investigating temperate^30, 32, 33^ and nucleus forming^19, 34^ phages. However, other Class 2 CRISPR effectors have encountered serious limitations in overcoming the diversity of genetic content encoded in phages^12, 19, 20, 35^. It remains unclear whether an RNA-targeting Cas13 can broadly protect bacteria from a wide range of dsDNA phages.

Here, we explored systematically the ability of a single Cas13a variant to restrict wide-ranging phage infections in *E.coli*. Phage infection assays show that LbuCas13a is a robust inhibitor of phage infections across *E. coli* phage phylogeny. Further, Cas13-mediated phage restriction is robust across a diversity of phage infection modalities, phage lifestyles, genes, and transcript features, enabling direct and specific phage interference. We demonstrate how Cas13’s potent, broad-spectrum antiviral activity can be employed as a sequence-specific counterselection system suitable for recovering phage variants with edits as minimal as a single codon. These results highlight the extraordinary exposure and vulnerability of phage RNA molecules during phage infection and provide a robust, generalizable strategy for phage genome engineering.

## Results

### Cas13 homologs sparsely populate bacteria phyla

Phages encode diverse anti-defense strategies against the bacterial defense systems they are likely to encounter^13, 32^, which in turn can render those systems ineffective for either phage immunity or phage engineering. To determine whether Cas13 might be useful as both a broad-spectrum phage defense and a phage genome editing tool, we began by investigating the distribution of Cas13 effectors across bacterial phyla. We performed a bioinformatic search for Cas13 proteins across NCBI and Genome Taxonomy Database (GTDB) genomes, culminating in a non-redundant set of 224 Cas13 protein sequences (Fig. 1; Supplementary Fig. 1). Consistent with prior classification efforts, Cas13 subtypes cluster into four clades 13a-d. We found Cas13b to be the most widespread, yet predominantly found within *Bacteroidota*. In contrast, Cas13c and Cas13d subtypes appeared least common, primarily found in *Fusobacteriota* and *Bacillota* (formerly *Firmicutes*), respectively. We found Cas13a to be phylogenetically more widely dispersed, although relatively limited in the total number of homologs, spread across *Pseudomondota* (previously *Proteobacteria*), *Bacillota*, *Bacteroidota*, and *Fusobacteriota*.

**Fig. 1.**
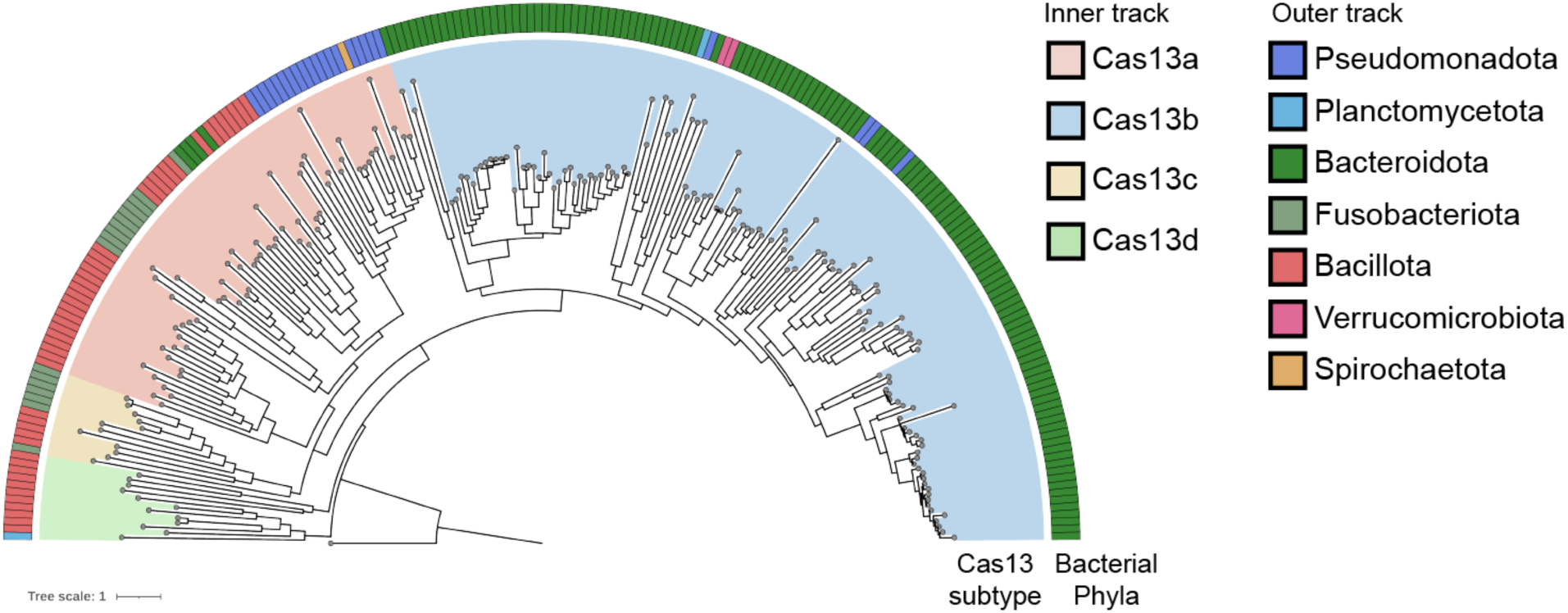
Maximum-likelihood phylogeny of Cas13 proteins and their distribution across the bacterial tree of life. The four known subtypes, Cas13a-d, each form their clade. A *Vibrio cholerae* Cas9 (UIO88932.1) was used as the outgroup. The microbial taxa that encode these Cas13s are denoted in the color bar.

Our results are consistent with prior CRISPR-search endeavors, suggesting that Cas13 effectors are some of the rarest Cas proteins currently identified^2^. Although RNA-targeting Type-III CRISPR-Cas systems are relatively abundant in bacterial phyla^2^, we wondered whether the sparse occurrence of Cas13 effectors means that general (ex. RNA-repair or RNA-modification) or specific (ex. anti-CRISPR^32^) resistance mechanisms to Cas13-activity are relatively rare as well.

### Cas13a is a more potent anti-phage effector than Cas13d

Two parsimonious explanations for the phylogenetic distribution of Cas13 effectors are that (1) Cas13 effectors are relatively ineffective anti-phage systems, limiting their phylogenetic spread from evolutionary pressure or (2) Cas13 effectors are potent anti-phage systems, but the fitness cost of their abortive-infection- (abi-) like effects^30, 33^ is selected against. To explore these possibilities, we tested the anti-phage activity of the most- and least-widely dispersed Cas13 effectors based on our analysis of bacterial phylogeny, Cas13a and Cas13d, respectively (Fig. 1). We selected LbuCas13a from *Leptotrichia buccalis* and RfxCas13d from *Ruminococcus flavefaciens* due to their extensive prior biochemical characterization^29, 36–39^. Notably, neither Cas13 variant has been investigated for antiviral activity. While a Cas13a ortholog from *Listeria seeligeri* has been used to restrict temperate and nucleus-forming phages^19, 30, 32, 34^, LbuCas13a comes from a phylogenetically distinct sub-clade of Cas13a effectors (Supplementary Fig. 1).

To develop a *E. coli* phage-challenge assay for LbuCas13a and RfxCas13d, we created “all-in-one” plasmids for inducible expression of *cas13* using anhydrotetracycline (aTc) alongside a constitutively expressed crRNA (DR-spacer) (Fig. 2a, b). During phage infection, phage RNAs are transcribed including a crRNA-targeted transcript (orange, Fig. 2a). Upon recognition, Cas13 activates HEPN-mediated RNA cleavage, although the extent of trans-cleavage may be reduced for Cas13d relative to Cas13a^37^. Depending on the extent of Cas13-mediated RNA cleavage, phage-encoded Cas13-resistance, protospacer mutation rate, and phage-encoded function containing the protospacer, phage may overcome the resulting general transcript degradation.

**Fig 2.**
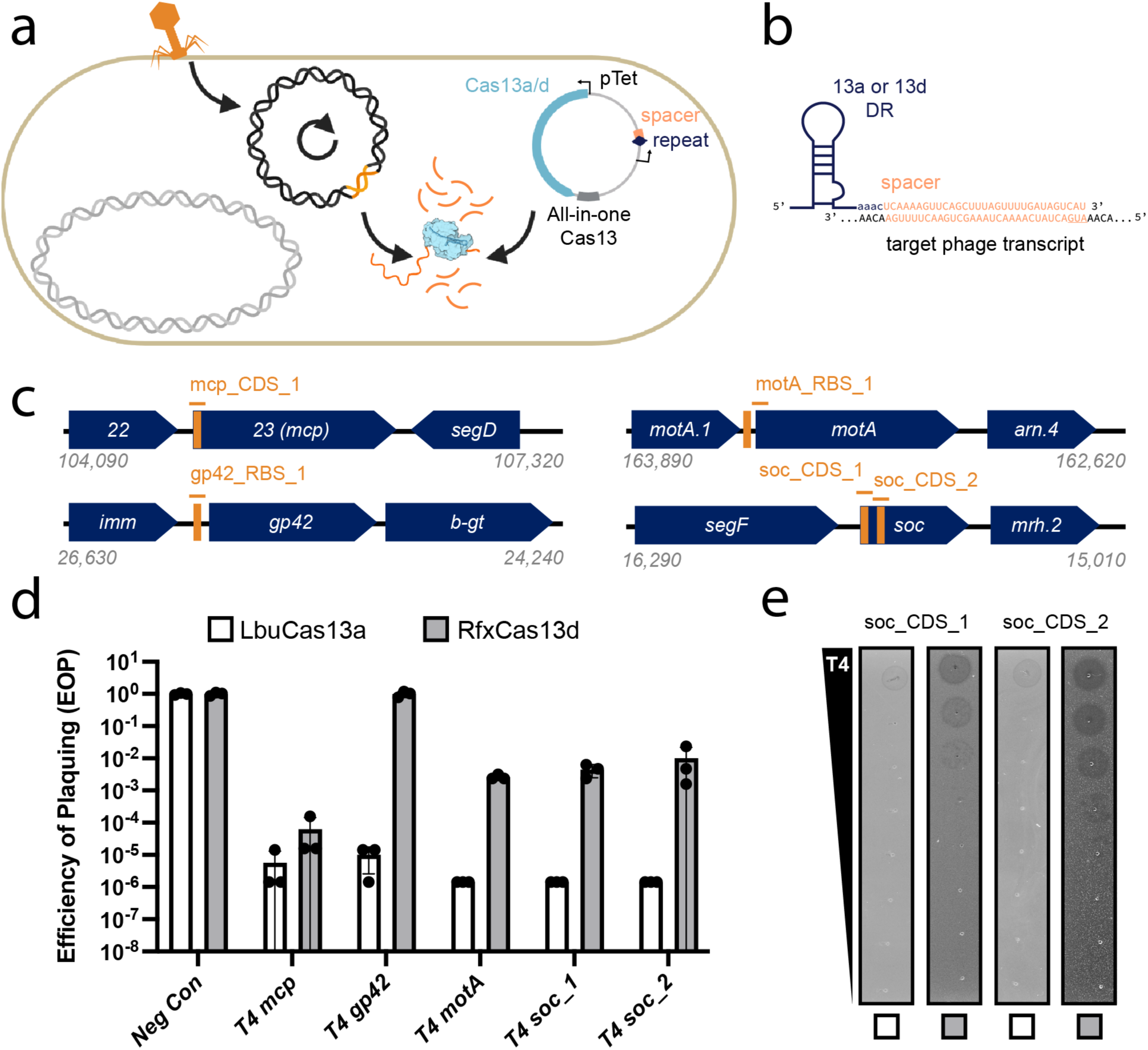
Comparison of Cas13a and Cas13d in *E. coli* phage-challenge assays with lytic phage T4. **(a)** Experimental architecture of Cas13 phage defense. Cas13 is expressed under anhydrotetracycline (aTc) control alongside a crRNA. During phage infection, Cas13 unleashes toxic cis- and trans-cleavage if Cas13 detects its crRNA target. **(b)** crRNA architecture employed in this study. **(c)** Overview of T4 genes and transcript locations targeted by Cas13 in T4 phage challenge experiments. Approximate gene architecture is shown in forward orientation. crRNA locations are highlighted in orange. **(d)** T4 phage infection in bacteria expressing phage-targeting crRNA and either LbuCas13a or RfxCas13d. EOP values represent the average of three biological replicates for a single crRNA. **(e)** T4 phage plaque assays comparing efficacy and toxicity of Cas13a and Cas13d. A representative plaque assay from three biological replicates is shown.

To test the phage-restriction capacity of LbuCas13a and RfxCas13d outside of their native context, we individually targeted a small panel of genes in phage T4. Phage T4 is a classical virulent dsDNA phage with a 170kb genome and well-characterized genetic content^40, 41^. From the perspective of phage genome editing, T4 represents an empirical challenge, displaying considerable variability in Cas-restriction efficacy for Cas9 and Cas12a, owing in part to modified glucosyl-5-hydroxymethylcytosine nucleotides^26, 27, 35^ and endogenous DNA-repair mechanisms^20^. For these reasons, we hypothesized RNA targeting could be a superior strategy to inhibit T4 and related phages. We designed a panel of Cas13 crRNAs targeting T4 transcripts with diverse design criteria (Fig. 2c)^40^. Targeted regions of T4^40^ RNA sequences included essential-genes (*mcp*, *motA*), a conditionally-essential gene (*gp42*), a non-essential gene (*soc*), an early-infection gene (*motA*), a middle-infection gene (*gp42*), late-infection genes (*mcp*, *soc*), encompassing regions early in coding sequences (CDSs) (*mcp*, *soc*), middle in CDS (*soc*) and untranslated regions around the ribosome binding site (RBS) (*gp42*, *motA*) (Supplementary Table 1). Broadly, this panel of crRNAs represents a systematic exploration of Cas13 targeting the diversity of feature types present in a phage transcriptome.

During phage infection experiments, we remarkably observed robust phage knockdown for all crRNAs tested using LbuCas13a (Fig. 2d). Independent of gene essentiality, timing of expression or position on transcript, we found that crRNA-guided LbuCas13a could restrict phage T4 over 100,000X with no substantial escape mutants observed (Supplementary Fig. 8). In contrast, crRNA-guided RfxCas13d exhibited highly variable and less-efficient phage restriction. Further, RfxCas13d exhibited phage-independent *E. coli* growth inhibition during RfxCas13d expression (Fig. 2e) and also observed a high degree of phage escape for RfxCas13d relative to LbuCas13a (Fig. 2e). These results suggest that LbuCas13a is a remarkably potent restrictor of phage T4 relative to other CRISPR-Cas systems^20, 26, 27, 35^.

### Cas13a confers resistance across E. coli phage phylogeny

To the best of our knowledge, no single Cas effector (or antiviral defense protein) has been shown to confer broad-spectrum phage-resistance when pressured against diverse dsDNA phages. To uncover the phage-phylogenetic limits of Cas13a anti-phage activity, we challenged *E. coli* expressing LbuCas13a with a phylogenetically diverse panel of dsDNA *E. coli* phages. To generate a representative sampling of *E. coli* phages, we constructed a protein-sharing network from 2307 phage genomes visualizing the relatedness of currently known *E. coli* phages (Fig. 3a). From this network, we assembled a panel of eight dsDNA *E. coli* phages scattered across coliphage phylogeny (Fig. 3a, b, Table 1). This panel includes both model *E. coli* phages (T4, T5, T7, and λ) and non-model *E. coli* phages (EdH4, MM02, N4, and SUSP1). With the sole exception of phages T4 and MM02, these phages bore minimal nucleotide sequence similarity to each other (Fig. 3a, b). Furthermore, these phages have diverse lifestyles and reflect a realistic model-sampling of the genetic diversity found among known dsDNA phages. Only one of these phages is temperate (λ) while the remaining seven are obligately lytic. They comprise diverse lifestyles including documented superspreader^42^, DNA compartmentalization^16^ and pseudolysogeny^43^ phenotypes. In aggregate, these phages not only represent genotypic diversity but also encompass a mixture of host-takeover strategies, modes of entry, and degrees of prior characterization.

**Fig 3.**
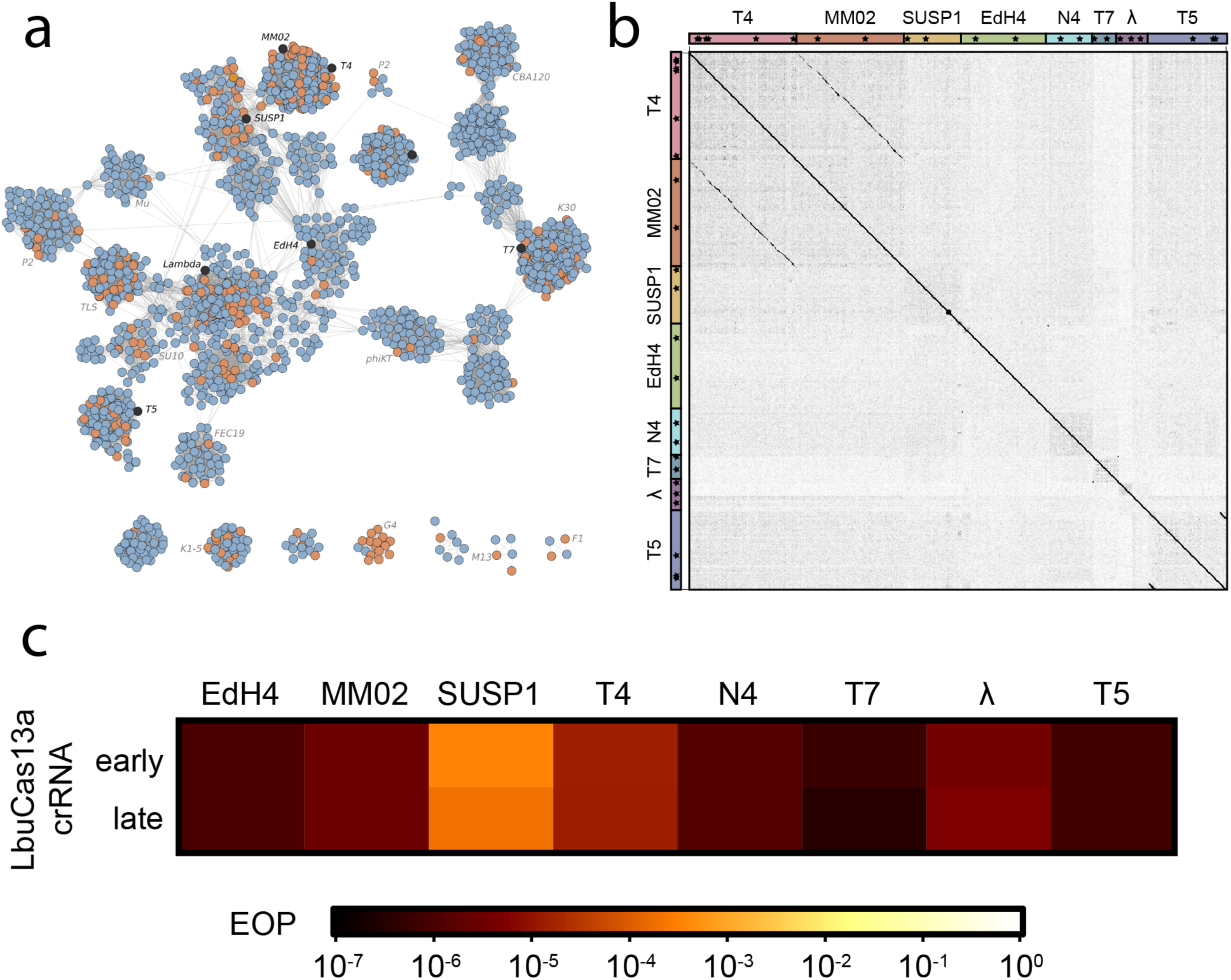
Comparison of LbuCas13a anti-phage activity across dsDNA *E. coli* phage phylogeny. **(a)** Network graph representation of *E. coli* phages and their relatives. Nodes represent phage genomes that are connected by edges if they share significant similarity as determined by vContact2^65^ (protein similarity). Nodes are shaded red if they are classified as an *E. coli* phage and blue if they only share similarity. Nodes are shaded black if they were assessed for sensitivity to LbuCas13a. **(b)** Gepard^66^ dot plot comparing the average nucleotide identity of phage genomes and the location of crRNAs used in this study. Higher regions of similarity are shown with darker shade. Phage genomes are concatenated and annotated on the axes. **(c)** Efficiency of plaquing (EOP) experiments for Cas13a designed to target an early- or late-transcript. EOP values represent the average of three biological replicates for a single crRNA.

**Table 1.** Summary of crRNAs Used in Cas13a Phage Restriction Assays.

| Phage | Phage Family | Genes Targeted | crRNA Success Rate (>1000X Restriction) | Reference genome |
| --- | --- | --- | --- | --- |
| $\lambda$ | *Siphoviridae*<br>(*Lambdavirus*) | *cro*, *ea47*, *mcp* | 3/3 (100%) | J02459.1 |
| EdH4 | *Myoviridae*<br>(*Vequintavirinae*) | *dnap*, *mcp* | 2/2 (100%) | MK327930.1 |
| MM02 | *Myoviridae*<br>(*Tevenvirinae*) | *dnap*, *mcp* | 2/2 (100%) | MK373784.1 |
| N4 | *Schitoviridae*<br>(*Enquatrovirinae*) | *dnap*, *mcp* | 2/2 (100%) | NC_008720.1 |
| SUSP1 | *Myoviridae*<br>(*Ounavirinae*) | *dnap*, *mcp* | 2/2 (100%) | NC_028808.2 |
| T4 | *Myoviridae*<br>(*Tevenvirinae*) | *dnap*, *gp42*,<br>*mcp*, *soc* | 5/5 (100%) | NC_000866.4 |
| T5 | *Demerecviridae*<br>(*Markadamsvirinae*<br>) | *dnap* (eLbu),<br>*gp150*, *mcp* | 3/3 (100%) | NC_005859.1 |
| T7 | *Autographiviridae*<br>*(Studiervirinae)* | *rnap, mcp* | 2/2 (100%) | NC_001604.1 |

For each phage, we designed a pair of Cas13a crRNAs targeting either a putative early gene (DNA polymerase (*dnap*), RNA polymerase (*rnap* (T7)), a lytic regulator (*cro*) or a putative late gene (major capsid protein (*mcp*)). An overview of Cas13-mediated phage restriction can be found in Table 1, diversity of crRNAs tested shown in Supplementary Fig. 2, and a by-phage summary of results shown in Supplementary Figs. 3-10. In aggregate, we observed substantial anti-phage activity for all 16 guides across the 8 phages tested (Fig. 3c, Table 1). Most guides reduced phage infectivity 10^5^-10^6^- fold, with the rare observation of escape mutants. Across this entire study, only a pair of escape mutants at 10^-4^ percent abundance were observed for T7 *mcp* targeting experiments (Supplementary Fig. 10). We observed a single guide (targeting T5 *dnap*) to yield general toxicity and growth inhibition during LbuCas13a induction. This constraint required us to perform assays in the absence of induction, achieving a mere 10^2^-fold knockdown (Supplementary Fig. 9). However, employing a reduced-toxicity LbuCas13a mutant^39^, we observed phage restriction at 10^6^-fold, suggesting that the subpar performance was due to baseline toxicity, rather than target toxicity.

Interestingly SUSP1 consistently displayed a small degree of resistance to Cas13a (Fig. 3c). Both early- and late-transcript targeting guides only decreased phage infectivity ∼5000-fold compared to all other phages showing 10^5^-10^6^-fold infectivity reduction. We further investigated the efficacy of SUSP1-targeting crRNAs in a plate-reader assay at a wide-range of multiplicities of infection (MOIs) (Supplementary Fig. 11). Compared to a non-targeting crRNA control, we found that both SUSP1*dnap-* and SUSP1*mcp*-targeting guides conferred phage resistance at all MOIs tested, including MOIs >10. These results indicate that Cas13a provided substantial protection against SUSP1 infection at both single-cell and population-levels^30^. Overall, LbuCas13a is capable of anti-phage activity with no apparent limits across phage phylogeny.

### LbuCas13a mediated phage restriction is independent of phage gene essentiality

We wondered whether the essentiality of the Cas13-targeted phage gene matters for Cas13a-mediated phage restriction. Prior work suggests that Cas13a primarily imparts phage defense through RNA trans-cleavage activity, but these observations derive from either temperate phages^30, 32^ or from targeting essential genes^19, 34^. However, for the majority of phages probed here, gene essentiality is poorly annotated. Therefore, we extended our study to crRNAs that target known non-essential genes of temperate phage λ (*ea47*)^44^, virulent phage T4 (*soc*)^40^, and virulent phage T5 (*gp150*)^45^. Across non-essential phage genes, we found Cas13a-mediated restriction to be as effective as when targeting essential phage genes (Fig. 4), indicating that the primary counterselection pressure does not depend on the essentiality of the crRNA target.

**Fig. 4.**
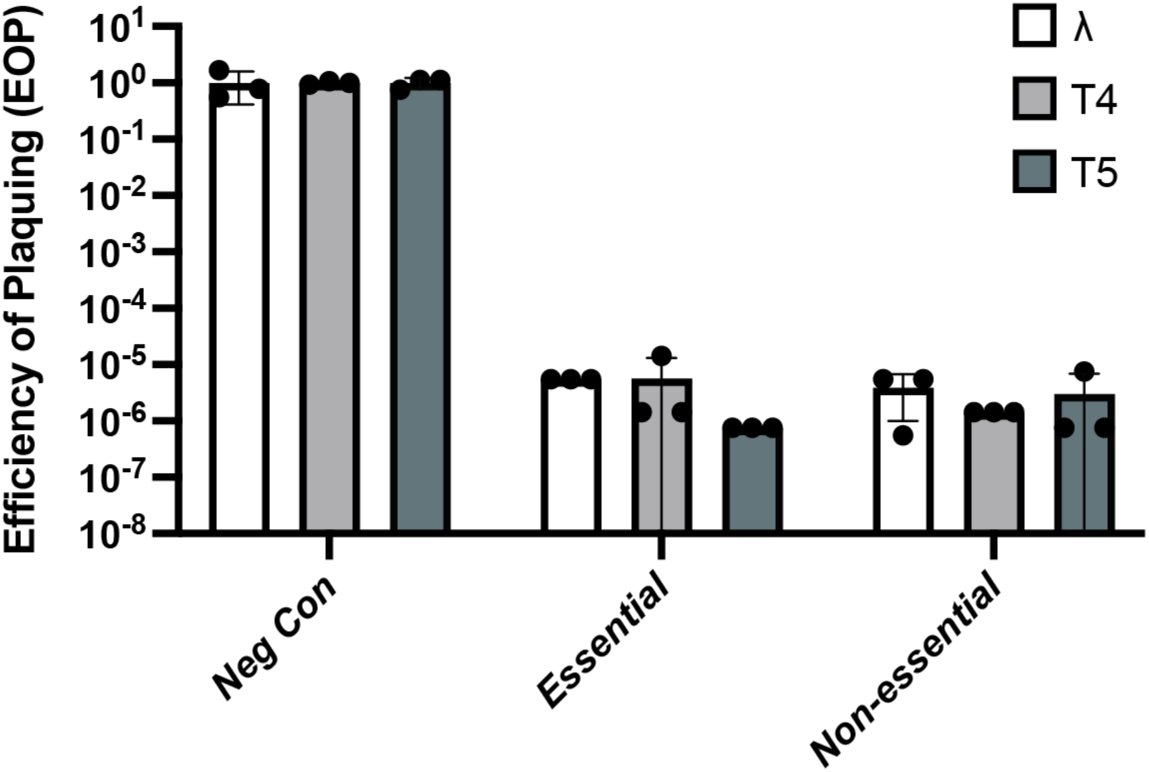
Anti-phage activity of Cas13a targeting essential versus non-essential phage genes. Different EOP experiments employing Cas13a crRNAs target either phage-essential or non-essential genes in diverse phages λ, T4, and T5; EOP values represent the average of three biological replicates for a single crRNA.

### A generalizable, markerless method for editing virulent phage genomes using Cas13a

The editing of virulent phage genomes has remained a major challenge for phage engineering, largely due to the lack of universally applicable genetic tools or reliance on a native CRISPR-Cas system^25, 26, 28, 46–50^. While the introduction of foreign gene content into phages is relatively straightforward to perform with homologous recombination (HR), ultimately the selection or screening for these rare recombinants is limiting even in well characterized phages^48^. Given that our LbuCas13a phage-restriction efficacy appears to have very little variability in terms of guide- (Fig. 2), target- (Figs. 3, 4), and phage-choice (Fig. 3), we suspected that Cas13a-mediated phage restriction would be an ideal tool for counterselection during phage genome editing. The high counter-selection stringency observed earlier in this study obviates the need for selection markers creating opportunities for multi-loci editing. Furthermore, the absence of PAM requirements for LbuCas13a targeting^29^ suggests that virtually any position within or nearby a phage transcript could be edited and selected through LbuCas13a counterselection.

We aimed to take advantage of these features by creating and enriching minimal edits that only Cas13a could easily select for^20, 26, 27, 35^, using T4 as a model virulent phage. We designed 6 mutants at either the non-essential *soc* gene or essential *dnap* using silent mutations, thus “re-coding” the target gene (Fig. 5). We designed these mutants to re-code only a single codon (*soc*-C, *dnap*-C), re-code the entire seed region (*soc-*S, *dnap*-S)^38^, or re-code the full target (*soc-*F, *dnap*-F) (Fig. 5acf). To facilitate homologous recombination-mediated edits, we flanked the intended mutation with 52 bp of native phage homology (Fig. 5a).

**Fig 5.**
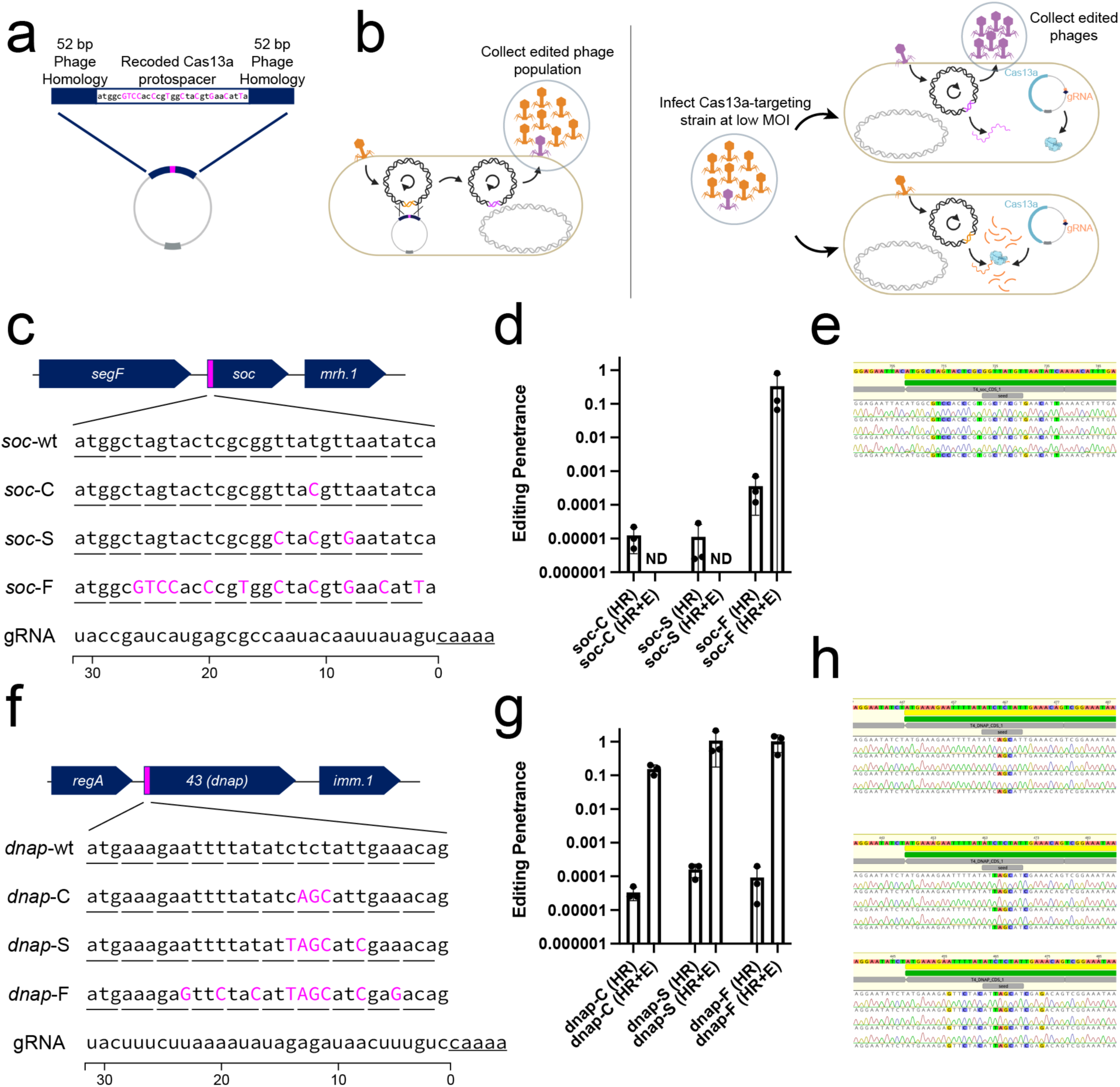
Editing of phage T4 in response to Cas13a counterselection. **(a)** Homologous recombination vector design consists of a re-coded Cas13a protospacer flanked by 52 bp of homology to the phage genome. **(b)** Overview of a simple 2-step editing process. Wildtype phage T4 infects homology vector-containing strain at a low MOI, yielding a mixed population of wt (orange) and edited (purple) phages (“HR”). This population is diluted and infects a LbuCas13a-expressing strain targeting the wt locus (10 nM aTc), enriching for edited phages relative to wt (“HR+E”). **(c)** Re-coding design for a T4 non-essential gene, *soc*, with introduced silent mutations shown in magenta. Three designs with differing mutations were tested (*soc*-C, *soc*-S, *soc*-F). **(d)** Survivor ratios from three biological replicates of the editing and enrichment process shown in (b) for *soc*-C, *soc*-S, *soc*-F. **(e)** Unbiased sequencing of T4 *soc* loci from individual plaques from three independent editing attempts. Deviations from wildtype are highlighted. **(f)** Re-coding design for a T4 essential gene, *dnap*, with introduced silent mutations shown in magenta. 3 designs with differing degrees of mutations were tested (*dnap*-C, *dnap*-S, *dnap*-F). **(g)** Survivor ratios from 3 biological replicates of the editing and enrichment process shown in (b) for *dnap*-C, *dnap*-S, *dnap*-F. **(h)** Unbiased, sequencing of T4 *dnap* loci from individual plaques from three independent editing attempts. Deviations from wildtype are highlighted. Sanger sequencing traces for all verified plaques including those shown in panels (e, h) can be found in Supplementary Figs. 16, 20).

In principle, edits in the phage genome introduced through homologous recombination can escape LbuCas13a targeting, while wildtype phage can not (Fig. 5b). To introduce and select for edits, we performed a simple, two-stage homologous recombination and enrichment process (Fig. 5b, Supplementary Fig. 12, Methods). Briefly, we employed two strains per edit: an editing strain containing a homologous recombination vector hosting a re-coded protospacer as well as 52 bp of phage homology (Fig. 5a,c,f) and a counterselection strain containing a LbuCas13a and crRNA expressing strain targeting either wt *soc* or *dnap* transcript (Fig. 2a,b). We first infected our editing strain with wildtype phage T4 at low MOI and collected the lysate, consisting of a mixture of wildtype and edited phages (“HR” phage lysate) (Fig. 5b, Supplementary Fig. 12a). Then we diluted this lysate, infected the counterselection strain at low MOI, and collected the resultant lysate (“HR+E” phage lysate) (Fig. 5b, Supplementary Fig. 12b). After each stage, lysates were collected and titered on the corresponding counterselection strain and a non-targeting control to assess editing penetrance.

For 4 of the 6 edits (*soc*-F, *dnap*-C, *dnap*-S, *dnap-*F), we observed plaques emerge at 0.1-0.001% abundance in the “HR” lysate (Supplementary Figs. 15, 17-19). After enrichment on the Cas13a counterselection strain, resistant plaques consisted of almost all of the phage population, suggesting high editing penetrance (Fig. 5d,g, Supplementary Figs. 15, 17-19). In contrast, lysates containing *soc*-C and *soc*-S mutations went to extinction following enrichment, suggesting that the *soc*-C and *soc*-S mutations were insufficient to evade Cas13a activation. Comparing the design of *soc*-C, *soc-*S, and *dnap*-C, multiple, contiguous mutations within the seed region appear necessary to evade Cas13a activation during phage infection. Potentially, one of the reasons we observed so few escape mutants is that multiple, contiguous mutations are needed to evade Cas13a activation.

To verify that Cas13a-resistant plaques were the result of intended edits, we performed unbiased PCRs at the wildtype locus for all editing attempts yielding plaques (Supplementary Figs. 15, 17-19). We analyzed plaques from 4 unique edits (*soc-F*, *dnap-C*, *dnap-S*, and *dnap-F*) and 12 independent editing processes, for a total of 36 plaques. Strikingly, we found all 36 analyzed plaques to have the intended mutation (Table 2, Supplementary Fig. 16, 20). This editing process represents a simple, straightforward route for enriching phage genome edits as small as one codon, as illustrated in the case of *dnap-C*.

**Table 2.** Summary of Cas13a-Mediated Phage Genome Editing.

| Edit Name | Phage | Edited Locus | Number of SNPs Introduced | Survivors Detected? | Plaques Screened | Mutant Success Rate |
| --- | --- | --- | --- | --- | --- | --- |
| *soc-C* | T4 | *soc* | 1 | No | N/A | N/A |
| *soc-S* | T4 | *soc* | 3 | No | N/A | N/A |
| *soc-F* | T4 | *soc* | 11 | Yes | 9 | 100% |
| *dnap-C* | T4 | *dnap* | 3 | Yes | 9 | 100% |
| *dnap-S* | T4 | *dnap* | 5 | Yes | 9 | 100% |
| *dnap-F* | T4 | *dnap* | 9 | Yes | 9 | 100% |

## Discussion

We have shown that LbuCas13a transcript targeting is a near-universal, programmable phage counterselection pressure that can be easily converted into a phage genome editing tool. Despite belonging to one of the rarest known CRISPR-Cas systems, we found LbuCas13a to be a potent RNA-guided anti-phage system. We challenged *E. coli* expressing Cas13a with eight diverse phages scattered across *E. coli* dsDNA phage phylogeny and found Cas13a to be effective at restricting all of them (>10^4^-fold). These results suggest that it is rare to encode mechanisms to broadly recover from or prevent RNA degradation in phages. Furthermore, we observed very high crRNA efficacy and consistency between these phages and designed targets. Cas13a anti-phage activity was consistent and effective across gene essentiality, gene expression timing, and target location. Due to the minimal constraints of crRNA design for LbuCas13a, we anticipate the primary constraint on crRNA design to be self-toxicity independent of the phage, a limitation that is uncommon and readily circumvented by designing an alternative crRNA. Due to the relative scarcity of Type-VI systems, we doubt many phages harbor Type-VI anti-CRISPR systems. Based on our observations, it appears that dsDNA phages are generally vulnerable to Cas13a targeting.

Leveraging the broad vulnerability of phages to Cas13a-targeting, we demonstrated how this robust counterselection could be employed to enrich markerless genome edits in *E. coli* phage. Most Cas-based counterselection methods show extensive crRNA or phage variability^25, 28, 34, 47^, rely on native CRISPR host-biology^46–49^, and/or yield a high rate of escape mutants^34, 46^. In contrast, we observed little variability in Cas13a counterselection efficacy across the eight phages and 21 crRNAs tested in this study. When applied to minimal, markerless genome editing that could only be selected using Cas13a, we measured a mutational penetrance of 100% - 36/36 plaques across 4 unique edits and 3 independent editing attempts (Fig. 5, Table 2). We anticipate this phage selection strategy can enrich nearly any viable edit at any phage locus in any phage whose host can harbor and express LbuCas13a.

Possibly, the highly potent anti-phage activity observed by Cas13a explains the relative scarcity of Type-VI CRISPR-Cas systems. All known Type-VI systems are thought to facilitate anti-phage activity through mechanisms similar to abortive infection ^30, 33^. Although the use of crRNA confers specificity for the activation of Cas13, we noticed substantial activation and toxicity in the absence of phage for RfxCas13d. Additionally, we observed substantial premature activation for one crRNA (targeting T5 *dnap*) with LbuCas13a that was remedied by using a less toxic variant of LbuCas13a^39^, highlighting this possible design constraint on crRNA designs. Nonetheless, the high reliability of crRNA efficacy we observe in tandem with flexible crRNA design afforded by Cas13a means that this occasional limitation is easily circumventable. Perhaps the genetic stability and performance of this phage-counterselection system would be more limited as it is applied in more diverse bacteria and their phages with higher mutation rates.

In some respects, the seemingly universal efficacy of Cas13a against phages is surprising. RNA-cleaving HEPN domains, such as those in Cas13a^6, 29^, are widely found across the tree of life including *E.coli* and related bacteria^51–53^. Although phages occasionally encode direct inhibitors of specific HEPN domains including Cas13a^32, 54^, it appears that general phage-encoded strategies for mitigating the toxic and anti-phage effects of HEPN-mediated RNA transcleavage are relatively limited. In contrast, phages encode a diversity of mechanisms to mitigate the effects of dsDNA cleavage including nuclease inhibitors^11, 13, 14, 55^, DNA modifications^15, 17^, DNA repair mechanisms^20, 21^, and nucleic acid compartmentalization^16, 18, 19^. This comparative vulnerability to degenerate RNA cleavage we observe for phages at large highlights the centrality of RNA for viral infection ^31^.

## Author Contributions

BAA and JAD conceived of the study and designed the experiments. BAA and TH performed informatic analysis. BAA conducted the experiments. BAA and BFC performed genetic design and molecular biology. BAA and VKM propagated phages. BAA wrote the initial manuscript. VKM, JFB, and RB contributed critical resources and advice. All authors contributed to the manuscript.

## Acknowledgments

We thank members of the Doudna lab, the Innovative Genomics Institute, and m-CAFEs SFA for helpful discussions, encouragement, and feedback. Original templates for LbuCas13a, eLbuCas13a, and RfxCas13d were kindly provided by Drs. Emeric Charles and David Savage. Phage SUSP1 was a generous gift from Dr. Sankar Adhya. We would also like to thank Dr. Hitomi Asahara, Dr. Eilleen Wagner, Angelica Lam, and Dr. Rohan Sachdeva for piloting custom full-plasmid sequencing and analysis pipelines that were instrumental to this study. The mature version of this pipeline is publicly accessible through the UC Berkeley Sequencing Facility. This project, BAA, and TH were supported by m-CAFEs Microbial Community Analysis & Functional Evaluation in Soils, a project led by Lawrence Berkeley National Laboratory based upon work supported by the U.S. Department of Energy, Office of Science, Office of Biological & Environmental Research. JAD is an Investigator of the Howard Hughes Medical Institute. Plasmids are available from Addgene (addgene.org) or by request.

## Methods

### Bacterial strains and growth conditions

Cultures of *E. coli* were grown in Lysogeny Broth (LB-Lennox) at 37°C, 250 rpm unless stated otherwise. When appropriate, 34 µg/mL chloramphenicol (+Ch) or 50 µg/mL kanamycin (+K) sulfate was supplemented to media. All bacterial strains were stored at - 80°C for long term storage in 25% sterile glycerol (Sigma). All cloning and strains were performed in dh10b genotype cells (NEB, Intact Genomics).

### Phage propagation and scaling

Phages were propagated through commonly used protocols in LB media or LB top agar overlays (0.7%) ^56^. Unless stated otherwise phages were propagated on *E. coli* BW25113 [*lacI*^+^*rrnB*_T14_ Δ*lacZ*_WJ16_ *hsdR*514 Δ*araBAD*_AH33_ Δ*rhaBAD*_LD78_ *rph-1* Δ*(araB–D)567* Δ*(rhaD–B)568* Δ*lacZ4787*(::*rrnB-3*) *hsdR514 rph-1*]. Phages N4, T4, T5, and T7 were scaled on *E. coli* BW25113 ^57^. Phage SUSP1 was a gift from Dr. Sankar Adhya and scaled on *E. coli* BW25113 ^42^. Phages EdH4 and MM02 were obtained from DSMZ culture collection and scaled on *E. coli* BW25113 (DSM 103295 and DSM 29475 respectively) ^58^. Phage λ cI857 *bor*::*kanR* was a gift from Dr. Drew Endy and scaled as described previously ^59^. All phages were titered through 2 µL spots of 10X serial dilution of phage in SM Buffer (Teknova) on *E. coli* BW25113 in a 0.7% top agar overlay.

### Plasmid construction

A description of all plasmids and oligonucleotides to build them can be found in Supplementary Table 2 and 3 respectively. All plasmids used in this study were verified using whole plasmid sequencing services offered by the UC Berkeley DNA Sequencing Facility. All plasmids were maintained as strains and maintained at -80°C in 25% glycerol (Sigma).

All-in-one LbuCas13a, eLbuCas13a, and RfxCas13d plasmids were designed to include a Cas13 effector under tetR-pTet control and a crRNA placeholder (DR-2xBsaI dropout site) under constitutive expression on a p15a-CmR backbone. LbuCas13a, eLbuCas13a, and RfxCas13d entry vectors were constructed through Gibson assembly (NEB, E2611L), yielding plasmids pBA559, pBA560, and pBA562 respectively. Assembly of pBA559, pBA560, and pBA562 used PCRs derived from pEJC 1.2 Lbu, pEJC 1.2 Lbu A12, and pEJC 1.5 CasRX vectors that were gifts from Drs. Emeric Charles and David Savage^39^. Gibson reactions were purified with DNA Clean & Concentrator-5 (Zymo Research) and electroporated into DH10b (NEB, Intact Genomics).

crRNA spacers were introduced to pBA559, pBA560, and pBA562 through BsaIHFv2 (NEB, R3733L) golden-gate assembly. For each spacer, golden-gate compatible template was created by 5’-phosphorylating with T4 PNK (NEB) and annealing of oligonucleotides. Golden Gate reactions for crRNA assembly were purified with DNA Clean & Concentrator-5 (Zymo Research), electroporated into DH10b (NEB, Intact Genomics), and plated on LB+Ch, 37°C.

HR donor vectors were assembled through BbsI (NEB, R0539L) golden gate assembly. pBA707 a BBR1-KanR vector with an RFP dropout cassette and 2xBbsI restriction was used as the backbone. 5’-phosphorylated with T4 PNK (NEB) and annealed oligonucleotides were used for UP-homology (oBA1761/oBA1762 or oBA1765/oBA1766), DN-homology (oBA1763/oBA1764 or oBA1767/oBA1768), and mutated protospacer (oBA1769/oBA1770, oBA1771/oBA1772, oBA1773/oBA1774, oBA1775/oBA1776, oBA1777/oBA1778, or oBA1779/oBA1780). Golden-gate reactions were purified with DNA Clean & Concentrator-5 (Zymo Research) and electroporated into DH10b (NEB, Intact Genomics) and plated on LB+K, 37°C. RFP-negative colonies were chosen for sequence-verification.

### crRNA design

A complete summary of the spacers used in this study can be found in Supplementary Table 1. We designed all Cas13a/d crRNAs as 31-nt spacers with no substantial bias against or towards any protospacer flanking sequence (PFS). Spacers were exclusively chosen to target predicted phage transcripts or a non-targeting control based on published genome sequences for phage λ (J02459.1), EdH4 (MK327930.1), MM02 (MK373784.1), N4 (NC_008720.1), SUSP1 (NC_028808.2), T4 (NC_000866.4), T5 (NC_005859.1), and T7 (NC_001604.1). Because DH10b harbors λ-like prophage, φ80lacZΔM15, spacers were designed to avoid similarity to the DH10b genome (NC_010473.1) ^60^. The majority of spacers were chosen to target the transcript of a target coding sequence (CDS). However, some spacers were chosen to target untranslated regions and are demarcated with “RBS”.

### Minimal edit homologous recombination donor vector design

HR donor vectors were designed with 52-nt of homology upstream (UP) and downstream (DN) of a targeted protospacer on the phage genome. To encode minimal edits, predicted codons were converted to silent mutations in a single codon (-C), seed region (-S), or full protospacer (-F) using a coding table for *E. coli*. When possible, codons were maximally altered and rare codons avoided to minimize non–Cas13-phenotypic consequence. The estimated seed region was estimated as previously observed *in vitro* ^38^.

### Efficiency of plaquing assays

Bacteriophage assays were conducted using a modified double agar overlay protocol. For each Cas13-crRNA-phage combination, a strain of dh10b (NEB, Intact Genomics) containing a Cas13-crRNA plasmid (Supplementary Table 2) was grown overnight 37°C, 250 rpm. To perform plaque assays, 100 µL of saturated overnight culture was mixed with molten LB Lennox top agar supplemented with appropriate inducer and antibiotics and decanted onto a corresponding LB Lennox Agar plate (to final overlay concentrations of 0.7% (w/v) agar, 5 nM anhydrotetracycline (aTc), and 34 µg/mL chloramphenicol). For all phage experiments in this study no supplementary CaCl_2_ or MgSO_4_ salts were added. For pBA675 and pBFC1053, toxicity was apparent at 5 nM aTc, so lower levels of aTc were used (0 aTc added and 1 nM aTc respectively). For pBA769 assays were performed at 10 nM aTc to achieve restriction against phage SUSP1. Overlays were left to dry for 15 minutes under microbiological flame. For each Cas13-crRNA-phage combination, 10X serial dilutions of the appropriate phage were performed in SM buffer (Teknova), and 2 µL of each dilution were spotted onto the top agar and allowed to dry for 10 minutes. Plaque assays were incubated at 37°C for 12-16 hours. After overnight incubation, plaques were scanned using a standard photo scanner and plaque forming units (PFUs) enumerated. In cases where individual PFUs were not enumerable, but clearings were observed at high phage concentrations, the most concentrated dilution at which no individual plaques were observed was counted as 1 PFU. Efficiency of plaquing (EOP) calculations for a given condition were performed by normalizing the mean of PFU for a condition to the mean PFU of a non-targeting control: mean(PFUcondition)/mean(PFUnegativecontrol). All plaque assays were performed in biological triplicate. Calculations were performed using GraphPad Prism.

### Liquid phage infection assays

Liquid phage experiments were performed in a Biotek plate reader at determined multiplicities of infection (MOIs). Briefly, for each Cas13-crRNA-phage combination, a strain of dh10b (NEB, Intact Genomics) containing a Cas13-crRNA plasmid (Supplementary Table 2) was grown overnight 37°C, 250 rpm. Strains were diluted in fresh media (LB + Ch + 10 nM aTc) to an OD of 0.04 and 200µL transferred to a 96-well plate (Corning 3904), achieving a final cell count of ∼8*10^6^ CFU/well. Appropriate phages were diluted in SM Buffer (Teknova) to a maximal titer of 10^11^ PFU/mL and 10X serially diluted 7 times. To begin phage infection, 1 µL of phage was added to achieve MOIs of 1.25*10^-6^ to 12.5. Infection was monitored in a Biotek Cytation 5 plate reader for 16 hours, 200 rpm shaking, 37°C with OD600 readings every 5 minutes. All infection assays were performed in biological triplicate beginning from 3 independent overnights of bacterial culture.

### T4 Phage genome editing experiments

A graphical overview of the phage genome editing experiments is shown in Supplementary Fig. 12. All assays were performed in biological triplicate beginning from 3 independent overnights of bacterial culture. All editing workflows occurred in parallel processes.

To create genome edited phage lysates, a phage editing strain consisting of dh10b (NEB, Intact Genomics) containing a homologous recombination vector (pBA787-pBA792) (Supplementary Table 2) was grown overnight in LB+K media at 37°C, 250 rpm. Strains were diluted into fresh media (LB+K) to an OD of 0.04 and 200µL transferred to a 96-well plate (Corning 3904), achieving a final cell count of ∼8*10^6^ CFU/well. Wildtype T4 was added to each well to achieve a MOI of 0.01 (∼8*10^4^ PFU of T4 phage). Infection was monitored in a Biotek Cytation 5 plate reader at 200 rpm shaking, 37°C with OD600 readings every 5 minutes. Infection was allowed to proceed until there was a visible population crash (∼4.5 hours). Lysates were transferred to a 96 well block (Greiner 780271-FD) and 1 drop of chloroform (Sigma) was added to lyse remaining bacteria. These lysates comprise a mixture of homologous recombination-edited T4 and wildtype T4 and comprised “HR” phage lysate. Blocks were covered with an aluminum seal (Corning 6570). “HR” phage lysates were stored at 4°C until use. “HR” phage lysates were titered before enrichment.

To enrich genome edited phage lysates, a phage counterselection strain consisting of dh10b (NEB, Intact Genomics) containing a “counterselection” Cas13a vector (pBA691 for *soc* or pBA778 for *dnap*) (Supplementary Table 2) was grown overnight in LB+Ch media at 37°C, 250 rpm. Strains were diluted into fresh media (LB+Ch+10 nM aTc) to an OD of 0.04 and 200 µL transferred to a 96-well plate (Corning 3904), achieving a final cell count of ∼8*10^6^ CFU/well. “HR phage lysate” was added to each well to achieve a MOI of 0.01 (∼8*10^4^ PFU of total phage titer). Infection was monitored in a Biotek Cytation 5 plate reader at 200 rpm shaking, 37°C with OD600 readings every 5 minutes. Infection was allowed to proceed until there was a visible population crash (∼7 hours). Lysates were transferred to a 96 well block (Greiner 780271-FD) and 1 drop of chloroform (Sigma) was added to lyse remaining bacteria. These lysates comprise an enriched mixture of homologous recombination-edited T4 and wildtype T4 and comprised “HR+E” phage lysate. Blocks were covered with an aluminum seal (Corning 6570). “HR+E” phage lysates were stored at 4°C until use.

### Determination of T4 phage genome editing penetrance

Determination of phage-editing penetrance was determined by plaque assay of “HR” and “HR+E” lysates on non-selective and wt-counterselective strains. For *soc* edits, 10 nM aTc induction was used for strains containing pBA620 as a negative control and pBA691 as a wt-counterselective Cas13 vector. For *dnap* edits, 5 nM aTc induction was used for strains containing pBA620 as a negative control and pBA778 as a wt-counterselective Cas13 vector. Penetrance was defined as PFU_counterselection_/PFU_negative_. Average penetrance was calculated across independent editing attempts. Penetrance calculations were performed in Graphpad Prism.

To confirm the genotype of edits, we performed unbiased PCRs followed by Sanger-sequencing. In all cases, unbiased PCRs were designed to amplify from 200 bp upstream and 200 bp downstream of the mutated protospacer, touching down outside the 52 bp flanking homologous recombination supplied from the editing vectors. PCRs were performed on 3 individual plaques from each “HR+E” lysate after plaquing on the counterselection strain. Plaques were picked into 50 µL SM Buffer (Teknova) and allowed to diffuse out of the plaque plug at 4°C overnight. To prepare for PCR and denature phage virions, 10 µL of these samples were transferred to PCR tubes and boiled at 100°C for 10 minutes. For *soc*-F edits, plaques were picked from “HR+E” *soc* lysate plaques on pBA691, amplified using oBA1783, oBA1784, and sequenced with oBA1783. For *dnap*-C, *dnap*-S, and *dnap*-F edits, plaques were picked from “HR+E” *soc* lysate plaques on pBA778, amplified using oBA1781, oBA1782, and sequenced with oBA1781. Additionally, the same procedure was performed on wildtype T4 at both *soc* and *dnap* loci.

### Cas13 phylogenetic tree

Cas13 annotated protein sequences were compiled from NCBI and were identified in GTDB r95 using custom cas13 HMMs. All sequences which did not contain two [R/Q/N/K/H/****H] sequence motifs were removed. CD-HIT v4.8.1 ^61^ was used to cluster sequences with a length cutoff of 0.9 and sequence similarity of 0.9. Sequences were then independently aligned using MUSCLE v3.8.31 and were manually trimmed in Geneious ^62, 63^. A maximum likelihood phylogenetic tree was built from the alignment using IQ-TREE v1.6.12 ^64^ with the following parameters -st AA -nt 48 -bb 1000 -m LG+G4+FO+I.

### Phage genome comparisons network

Protein-protein phage genome comparisons were performed with VConTACT2^65^ MCL clustering (*rel-mode Diamond*, *vcs-mode ClusterONE*) of the protein sequences of three *E. coli* phages EdH4 (MK327930.1) and vB_EcoM_MM02 (MK373784.1) together with those of the Prokaryotic Viral RefSeq 201 phage database. Produced viral clusters which did not contain *E. coli* phage nor shared an edge with a viral cluster containing any *E. coli* phage were removed, together with singletons, to simplify the network.

Average nucleotide identity phage genome comparisons were performed with Gepard^66^ using a word length of 10 bp. For source genomes, we used a concatenation of the eight phage genomes used in this study: T4 (NC_000866.4), MM02 (MK373784.1), SUSP1 (NC_028808.2), EdH4 (MK327930.1), N4 (NC_008720.1), T7 (NC_001604.1), λ (J02459.1), and T5 (NC_005859.1).

## References

1. Barrangou, R. et al. CRISPR provides acquired resistance against viruses in prokaryotes. Science 315, 1709–1712 (2007).

2. Makarova, K. S. et al. Evolutionary classification of CRISPR-Cas systems: a burst of class 2 and derived variants. Nat. Rev. Microbiol. 18, 67–83 (2020).

3. Brouns, S. J. J. et al. Small CRISPR RNAs guide antiviral defense in prokaryotes. Science 321, 960–964 (2008).

4. Garneau, J. E. et al. The CRISPR/Cas bacterial immune system cleaves bacteriophage and plasmid DNA. Nature 468, 67–71 (2010).

5. Zetsche, B. et al. Cpf1 is a single RNA-guided endonuclease of a class 2 CRISPR-Cas system. Cell 163, 759–771 (2015).

6. Abudayyeh, O. O. et al. C2c2 is a single-component programmable RNA-guided RNA-targeting CRISPR effector. Science 1–17 (2016).

7. Kazlauskiene, M., Kostiuk, G., Venclovas, Č., Tamulaitis, G. & Siksnys, V. A cyclic oligonucleotide signaling pathway in type III CRISPR-Cas systems. Science 357, 605–609 (2017).

8. Niewoehner, O. et al. Type III CRISPR-Cas systems produce cyclic oligoadenylate second messengers. Nature 548, 543–548 (2017).

9. Huang, C. J., Adler, B. A. & Doudna, J. A. A naturally DNase-free CRISPR-Cas12c enzyme silences gene expression. bioRxiv 2021.12.06.471469 (2021) doi:10.1101/2021.12.06.471469.

10. Knott, G. J. & Doudna, J. A. CRISPR-Cas guides the future of genetic engineering. Science 361, 866–869 (2018).

11. Bondy-Denomy, J., Pawluk, A., Maxwell, K. L. & Davidson, A. R. Bacteriophage genes that inactivate the CRISPR/Cas bacterial immune system. Nature 493, 429– 432 (2013).

12. Harrington, L. B. et al. A Broad-Spectrum Inhibitor of CRISPR-Cas9. Cell 170, 1224–1233.e15 (2017).

13. Pawluk, A., Davidson, A. R. & Maxwell, K. L. Anti-CRISPR: discovery, mechanism and function. Nat. Rev. Microbiol. 16, 12–17 (2018).

14. Shivram, H., Cress, B. F., Knott, G. J. & Doudna, J. A. Controlling and enhancing CRISPR systems. Nat. Chem. Biol. 17, 10–19 (2021).

15. Sinsheimer, R. L. Nucleotides from T2r+ Bacteriophage. Science 120, 551–553 (1954).

16. Lanni, Y. T. First-step-transfer deoxyribonucleic acid of bacteriophage T5. Bacteriol. Rev. 32, 227–242 (1968).

17. Weigele, P. & Raleigh, E. A. Biosynthesis and Function of Modified Bases in Bacteria and Their Viruses. Chem. Rev. 116, 12655–12687 (2016).

18. Chaikeeratisak, V. et al. Assembly of a nucleus-like structure during viral replication in bacteria. Science 355, 194–197 (2017).

19. Mendoza, S. D. et al. A bacteriophage nucleus-like compartment shields DNA from CRISPR nucleases. Nature 577, 244–248 (2020).

20. Wu, X., Zhu, J., Tao, P. & Rao, V. B. Bacteriophage T4 Escapes CRISPR Attack by Minihomology Recombination and Repair. MBio 12, e0136121 (2021).

21. Hossain, A. A., McGinn, J., Meeske, A. J., Modell, J. W. & Marraffini, L. A. Viral recombination systems limit CRISPR-Cas targeting through the generation of escape mutations. Cell Host Microbe 29, 1482–1495.e12 (2021).

22. Borges, A. L. et al. Bacteriophage Cooperation Suppresses CRISPR-Cas3 and Cas9 Immunity. Cell 174, 917–925.e10 (2018).

23. Landsberger, M. et al. Anti-CRISPR Phages Cooperate to Overcome CRISPR-Cas Immunity. Cell 174, 908–916.e12 (2018).

24. Varble, A. et al. Prophage integration into CRISPR loci enables evasion of antiviral immunity in Streptococcus pyogenes. Nature Microbiology 6, 1516–1525 (2021).

25. Martel, B. & Moineau, S. CRISPR-Cas: an efficient tool for genome engineering of virulent bacteriophages. Nucleic Acids Res. 42, 9504–9513 (2014).

26. Tao, P., Wu, X., Tang, W.-C., Zhu, J. & Rao, V. Engineering of Bacteriophage T4 Genome Using CRISPR-Cas9. ACS Synth. Biol. 6, 1952–1961 (2017).

27. Duong, M. M., Carmody, C. M., Ma, Q., Peters, J. E. & Nugen, S. R. Optimization of T4 phage engineering via CRISPR/Cas9. Sci. Rep. 10, 18229 (2020).

28. Wetzel, K. S. et al. CRISPY-BRED and CRISPY-BRIP: efficient bacteriophage engineering. Sci. Rep. 11, 6796 (2021).

29. East-Seletsky, A. et al. Two distinct RNase activities of CRISPR-C2c2 enable guide-RNA processing and RNA detection. Nature 538, 270–273 (2016).

30. Meeske, A. J., Nakandakari-Higa, S. & Marraffini, L. A. Cas13-induced cellular dormancy prevents the rise of CRISPR-resistant bacteriophage. Nature 570, 241– 245 (2019).

31. Baltimore, D. Expression of animal virus genomes. Bacteriol. Rev. 35, 235–241 (1971).

32. Meeske, A. J. et al. A phage-encoded anti-CRISPR enables complete evasion of type VI-A CRISPR-Cas immunity. Science 369, 54–59 (2020).

33. VanderWal, A. R., Park, J.-U., Polevoda, B., Kellogg, E. H. & O’Connell, M. R. CRISPR-Csx28 forms a Cas13b-activated membrane pore required for robust CRISPR-Cas adaptive immunity. bioRxiv 2021.11.02.466367 (2021) doi:10.1101/2021.11.02.466367.

34. Guan, J. et al. RNA targeting with CRISPR-Cas13a facilitates bacteriophage genome engineering. bioRxiv 2022.02.14.480438 (2022) doi:10.1101/2022.02.14.480438.

35. Bryson, A. L. et al. Covalent Modification of Bacteriophage T4 DNA Inhibits CRISPR-Cas9. MBio 6, e00648 (2015).

36. Liu, L. et al. The Molecular Architecture for RNA-Guided RNA Cleavage by Cas13a. Cell 170, 714–726.e10 (2017).

37. Konermann, S. et al. Transcriptome Engineering with RNA-Targeting Type VI-D CRISPR Effectors. Cell 173, 665–676.e14 (2018).

38. Tambe, A., East-Seletsky, A., Knott, G. J., Doudna, J. A. & O’Connell, M. R. RNA Binding and HEPN-Nuclease Activation Are Decoupled in CRISPR-Cas13a. Cell Rep. 24, 1025–1036 (2018).

39. Charles, E. J. et al. Engineering improved Cas13 effectors for targeted post-transcriptional regulation of gene expression. bioRxiv 2021.05.26.445687 (2021) doi:10.1101/2021.05.26.445687.

40. Miller Eric S. et al. Bacteriophage T4 Genome. Microbiol. Mol. Biol. Rev. 67, 86– 156 (2003).

41. Kutter, E. et al. From Host to Phage Metabolism: Hot Tales of Phage T4’s Takeover of E. coli. Viruses 10, (2018).

42. Keen, E. C. et al. Novel ‘Superspreader’ Bacteriophages Promote Horizontal Gene Transfer by Transformation. MBio 8, (2017).

43. Los, M., Wegrzyn, G. & Neubauer, P. A role for bacteriophage T4 rI gene function in the control of phage development during pseudolysogeny and in slowly growing host cells. Res. Microbiol. 154, 547–552 (2003).

44. Kellenberger, G., Zichichi, M. L. & Weigle, J. A mutation affecting the DNA content of bacteriophage lambda and its lysogenizing properties. J. Mol. Biol. 3, 399–408 (1961).

45. Huet, A., Duda, R. L., Hendrix, R. W., Boulanger, P. & Conway, J. F. Correct Assembly of the Bacteriophage T5 Procapsid Requires Both the Maturation Protease and the Portal Complex. J. Mol. Biol. 428, 165–181 (2016).

46. Kiro, R., Shitrit, D. & Qimron, U. Efficient engineering of a bacteriophage genome using the type I-E CRISPR-Cas system. RNA Biol. 11, 42–44 (2014).

47. Box, A. M., McGuffie, M. J., O’Hara, B. J. & Seed, K. D. Functional Analysis of Bacteriophage Immunity through a Type I-E CRISPR-Cas System in Vibrio cholerae and Its Application in Bacteriophage Genome Engineering. J. Bacteriol. 198, 578–590 (2016).

48. Bari, S. M. N., Walker, F. C., Cater, K., Aslan, B. & Hatoum-Aslan, A. Strategies for Editing Virulent Staphylococcal Phages Using CRISPR-Cas10. ACS Synth. Biol. 6, 2316–2325 (2017).

49. Mayo-Muñoz, D. et al. Anti-CRISPR-Based and CRISPR-Based Genome Editing of Sulfolobus islandicus Rod-Shaped Virus 2. Viruses 10, (2018).

50. Grigonyte, A. M. et al. Comparison of CRISPR and Marker-Based Methods for the Engineering of Phage T7. Viruses 12, (2020).

51. Anantharaman, V., Makarova, K. S., Burroughs, A. M., Koonin, E. V. & Aravind, L. Comprehensive analysis of the HEPN superfamily: identification of novel roles in intra-genomic conflicts, defense, pathogenesis and RNA processing. Biol. Direct 8, 15 (2013).

52. Doron, S. et al. Systematic discovery of antiphage defense systems in the microbial pangenome. Science 359, (2018).

53. Gao, L. et al. Diverse enzymatic activities mediate antiviral immunity in prokaryotes. Science 369, 1077–1084 (2020).

54. Otsuka, Y. & Yonesaki, T. Dmd of bacteriophage T4 functions as an antitoxin against Escherichia coli LsoA and RnlA toxins. Mol. Microbiol. 83, 669–681 (2012).

55. Krüger, D. H., Schroeder, C., Hansen, S. & Rosenthal, H. A. Active protection by bacteriophages T3 and T7 against E. coli B- and K-specific restriction of their DNA. Mol. Gen. Genet. 153, 99–106 (1977).

56. Kutter, E. & Sulakvelidze, A. Bacteriophages: Biology and Applications. (CRC Press, 2004).

57. Mutalik, V. K. et al. High-throughput mapping of the phage resistance landscape in E. coli. PLoS Biol. 18, e3000877 (2020).

58. Korf, I. H. E. et al. Still Something to Discover: Novel Insights intoEscherichia coli Phage Diversity and Taxonomy. Viruses 11, (2019).

59. Knott, G. J. et al. Structural basis for AcrVA4 inhibition of specific CRISPR-Cas12a. Elife 8, (2019).

60. Durfee, T. et al. The complete genome sequence of Escherichia coli DH10B: insights into the biology of a laboratory workhorse. J. Bacteriol. 190, 2597–2606 (2008).

61. Fu, L., Niu, B., Zhu, Z., Wu, S. & Li, W. CD-HIT: accelerated for clustering the next-generation sequencing data. Bioinformatics 28, 3150–3152 (2012).

62. Edgar, R. C. MUSCLE: multiple sequence alignment with high accuracy and high throughput. Nucleic Acids Res. 32, 1792–1797 (2004).

63. Kearse, M. et al. Geneious Basic: an integrated and extendable desktop software platform for the organization and analysis of sequence data. Bioinformatics 28, 1647–1649 (2012).

64. Nguyen, L.-T., Schmidt, H. A., von Haeseler, A. & Minh, B. Q. IQ-TREE: a fast and effective stochastic algorithm for estimating maximum-likelihood phylogenies. Mol. Biol. Evol. 32, 268–274 (2015).

65. Bin Jang, H., et al. Taxonomic assignment of uncultivated prokaryotic virus genomes is enabled by gene-sharing networks. Nat. Biotechnol. 37, 632–639 (2019).

66. Krumsiek, J., Arnold, R. & Rattei, T. Gepard: a rapid and sensitive tool for creating dotplots on genome scale. Bioinformatics 23, 1026–1028 (2007).

