## Supplementary Information for "RNA-targeting CRISPR-Cas13 Provides Broad-spectrum Phage Immunity"

**Supplementary Fig. 1. Unrooted graph of Cas13 phylogeny.**

**Supplementary Fig. 2. Phage-restriction LbuCas13a-crRNAs are scattered across diverse phage genomes.**

**Supplementary Fig. 3. Summary of results for LbuCas13a interference against phage EdH4.**

**Supplementary Fig. 4. Summary of results for LbuCas13a interference against phage MM02.**

**Supplementary Fig. 5. Summary of results for LbuCas13a interference against phage λ.**

**Supplementary Fig. 6. Summary of results for LbuCas13a interference against phage N4.**

**Supplementary Fig. 7. Summary of results for LbuCas13a interference against phage SUSP1.**

**Supplementary Fig. 8. Summary of results for LbuCas13a and RfxCas13d interference against phage T4.**

**Supplementary Fig. 9. Summary of results for LbuCas13a interference against phage T5.**

**Supplementary Fig. 10. Summary of results for LbuCas13a interference against phage T7.**

**Supplementary Fig. 11. SUSP1 displays sensitivity to LbuCas13a targeting during liquid infection.**

**Supplementary Fig. 12. Method overview for Cas13a Phage Editing.**

**Supplementary Fig. 13. Overview of phenotypic results for editing attempt *soc*-C.**

**Supplementary Fig. 14. Overview of phenotypic results for editing attempt *soc*-S.**

**Supplementary Fig. 15. Overview of phenotypic results for editing attempt *soc*-F.**

**Supplementary Fig. 16. Overview of genotyping results for *soc* editing attempts.**

**Supplementary Fig. 17. Overview of phenotypic results for editing attempt *dnap*-C.**

**Supplementary Fig. 18. Overview of phenotypic results for editing attempt *dnap*-S.**

**Supplementary Fig. 19. Overview of phenotypic results for editing attempt *dnap*-F.**

**Supplementary Fig. 20. Overview of genotyping results for *dnap* editing attempts.**

**Supplementary Table 1 - Summary of crRNA Design**

**Supplementary Table 2 - Plasmids Used in This Study**

**Supplementary Table 3 - Oligonucleotides Used in This Study**

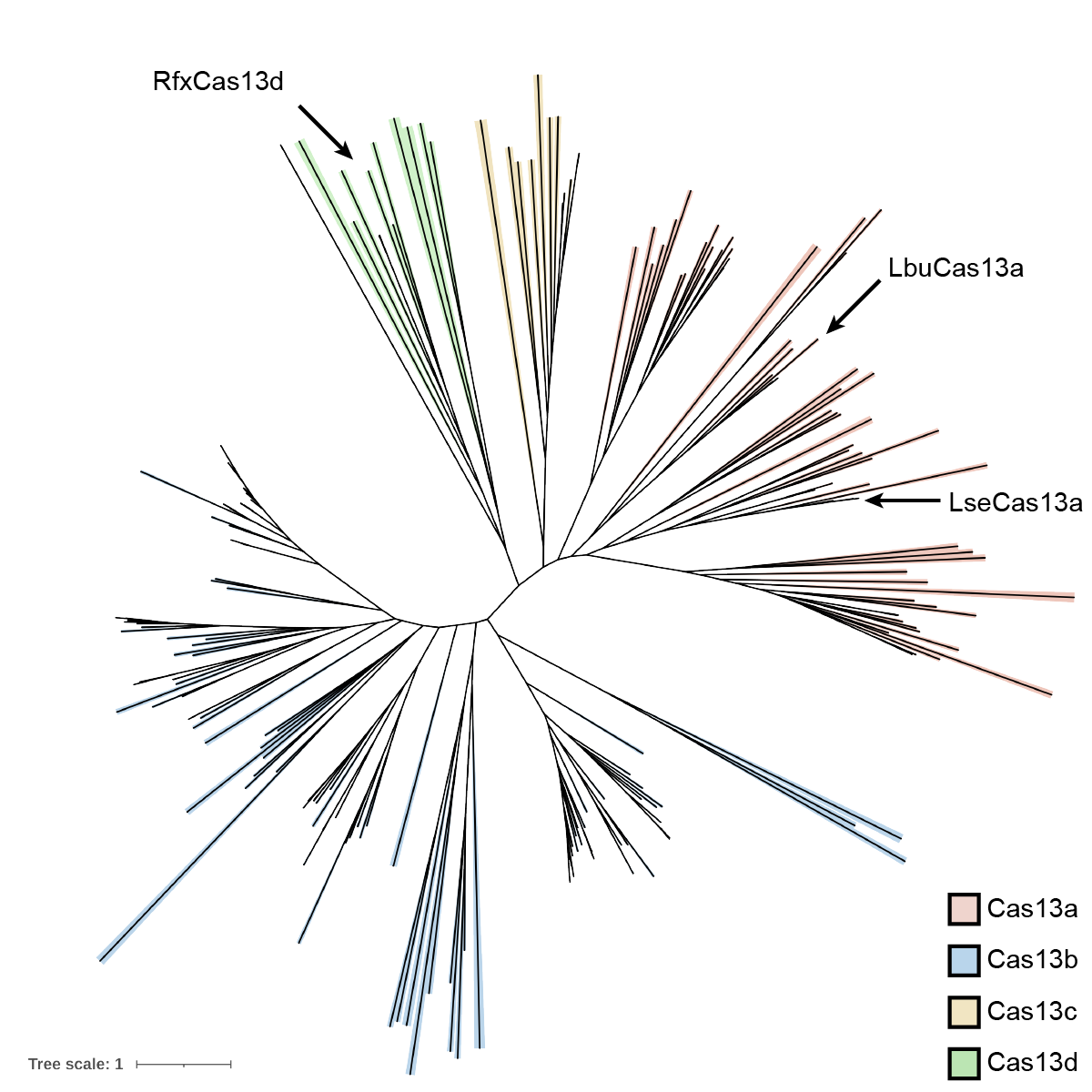

**Supplementary Fig. 1. Unrooted graph of Cas13 phylogeny.** Unrooted tree of Cas13 effectors colored by subtype. Cas13 effectors from this study (LbuCas13a, RfxCas13d) and prior studies (LseCas13a) investigating anti-phage activity are shown.

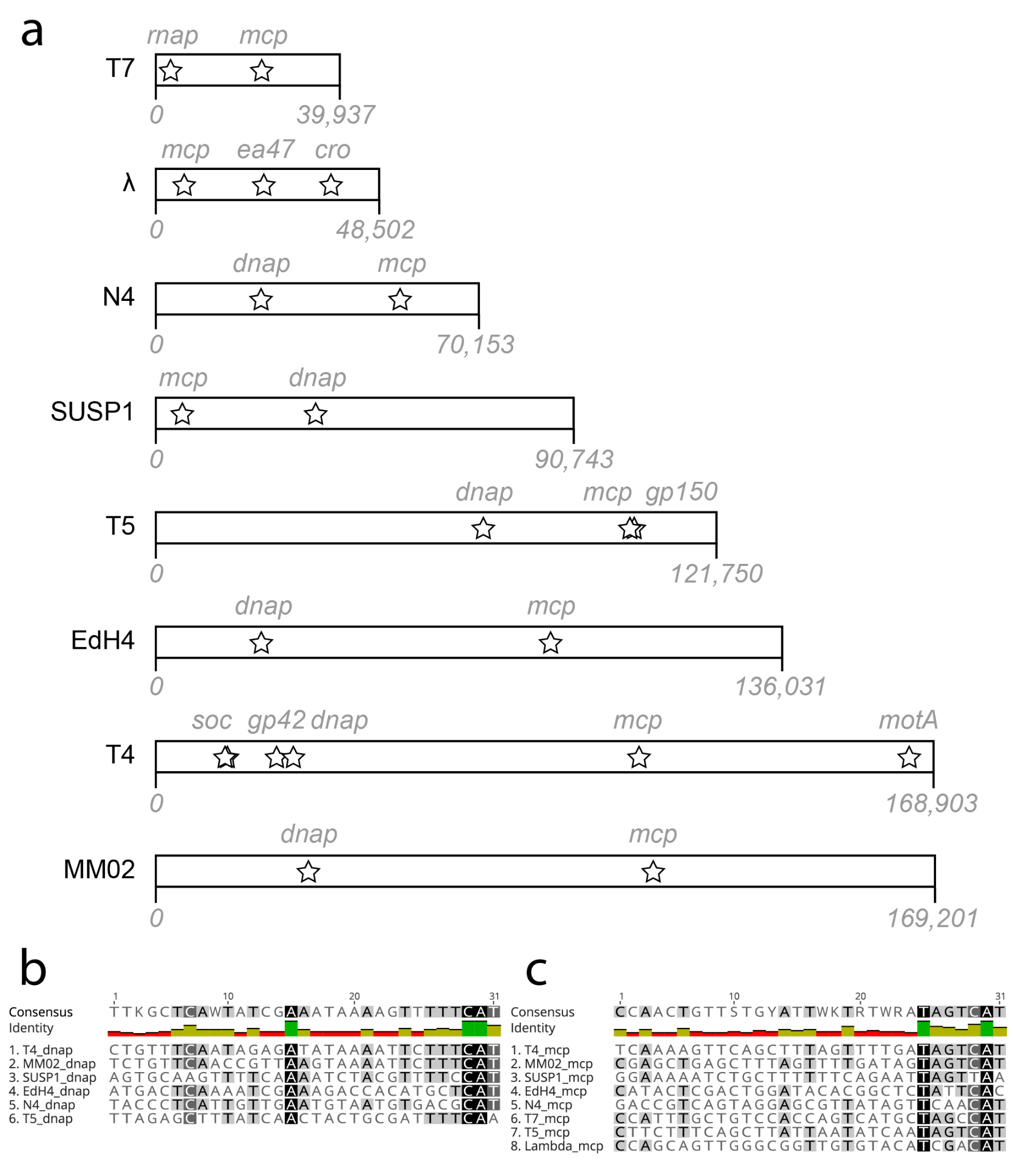

**Supplementary Fig. 2. Phage-restriction LbuCas13a-crRNAs are scattered across diverse phage genomes.** **(a)** Summary of phage’s genomes used in this study and locations of LbuCas13a crRNA targets (stars) mapped to scale. Target transcript’s closest gene is annotated. **(b)** Only distant similarity observed between spacers targeting the same gene, *dnap* across the phages investigated in this study. **(c)** Only distant similarity observed between spacers targeting the same gene, *mcp* across the phages investigated in this study.

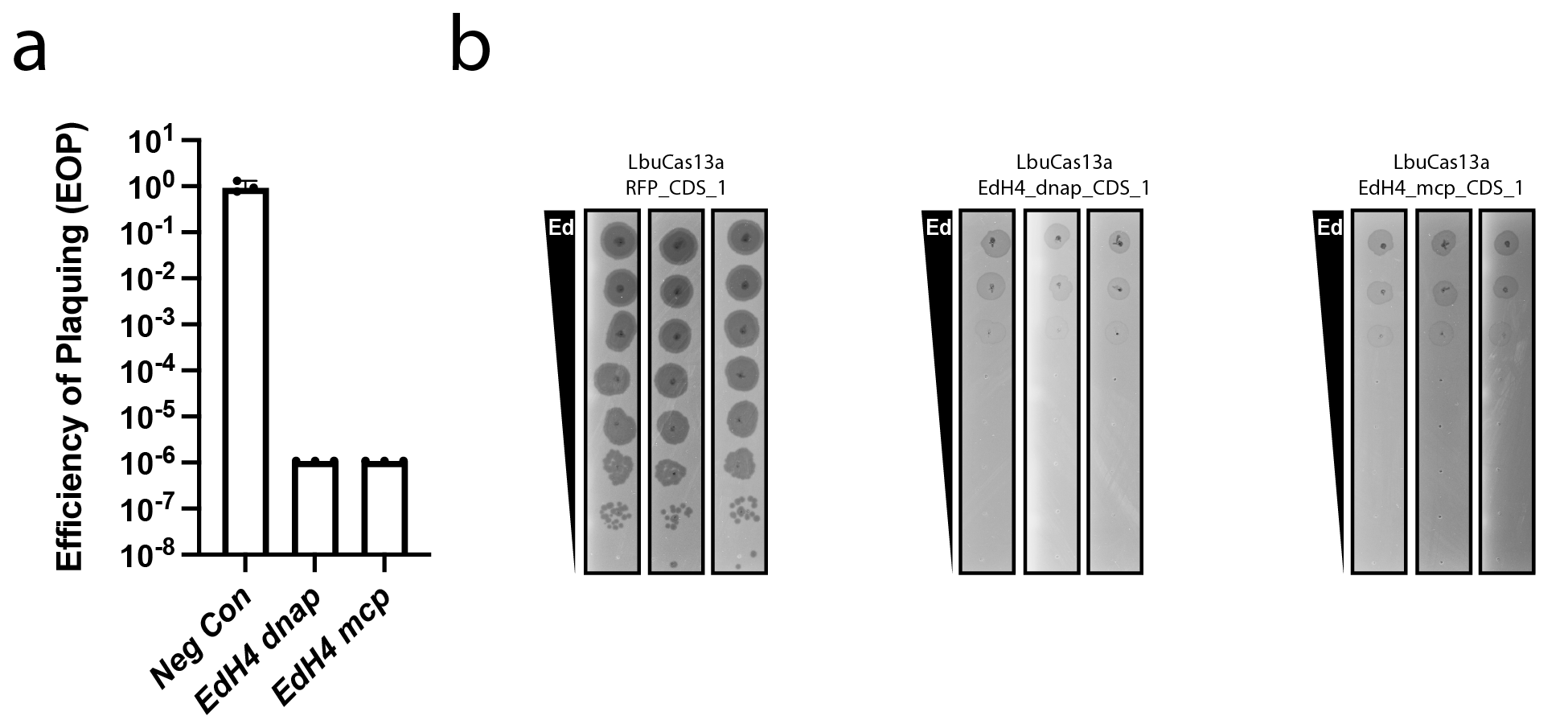

**Supplementary Fig. 3. Summary of results for LbuCas13a interference against phage EdH4.** **(a)** EOP estimation and **(b)** plaque assay results for EdH4 infection against all crRNAs tested at 5 nM aTc induction of LbuCas13a. For EOP graphs, results are represented as mean±std. All assays were performed in biological triplicate.

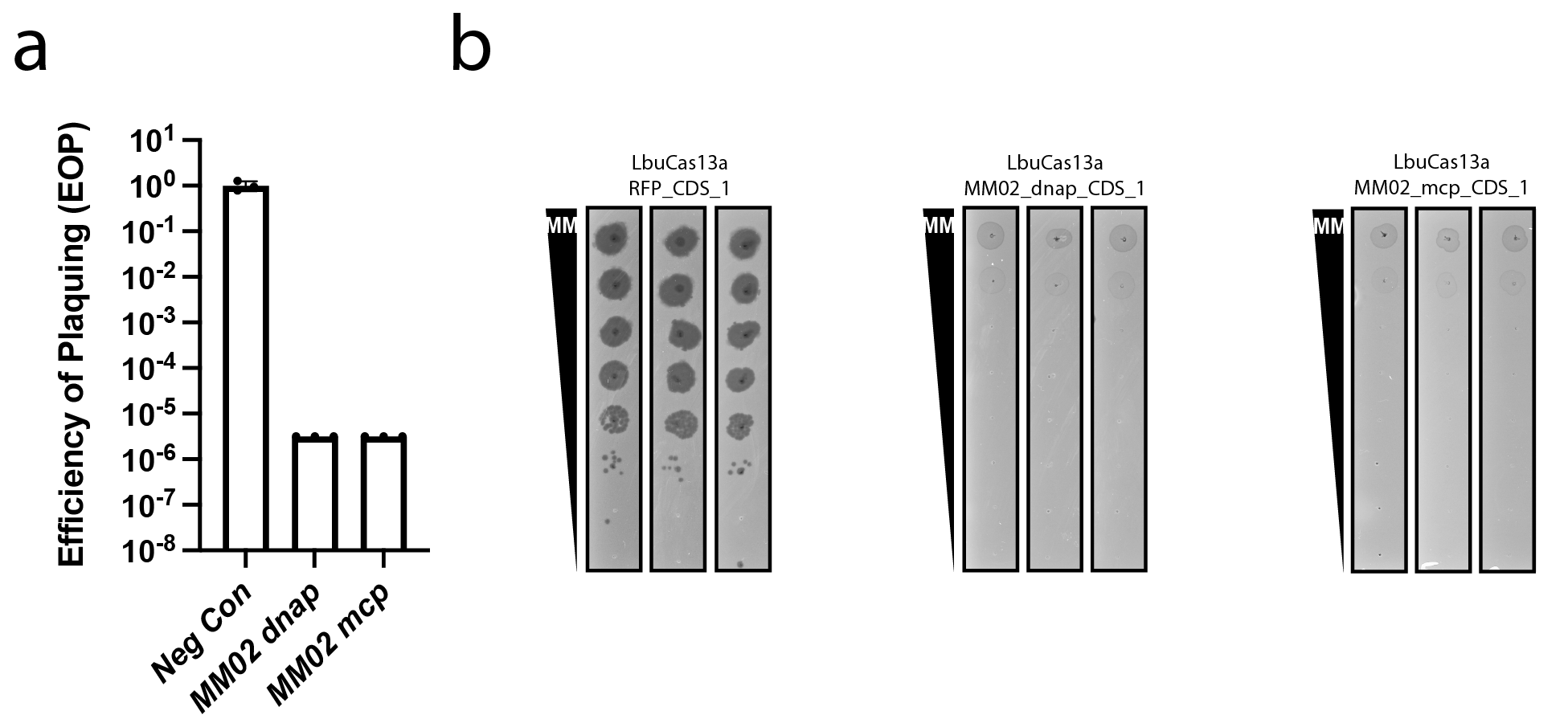

**Supplementary Fig. 4. Summary of results for LbuCas13a interference against phage MM02.** **(a)** EOP estimation and **(b)** plaque assay results for MM02 infection against all crRNAs tested at 5 nM aTc induction of LbuCas13a. For EOP graphs, results are represented as mean±std. All assays were performed in biological triplicate.

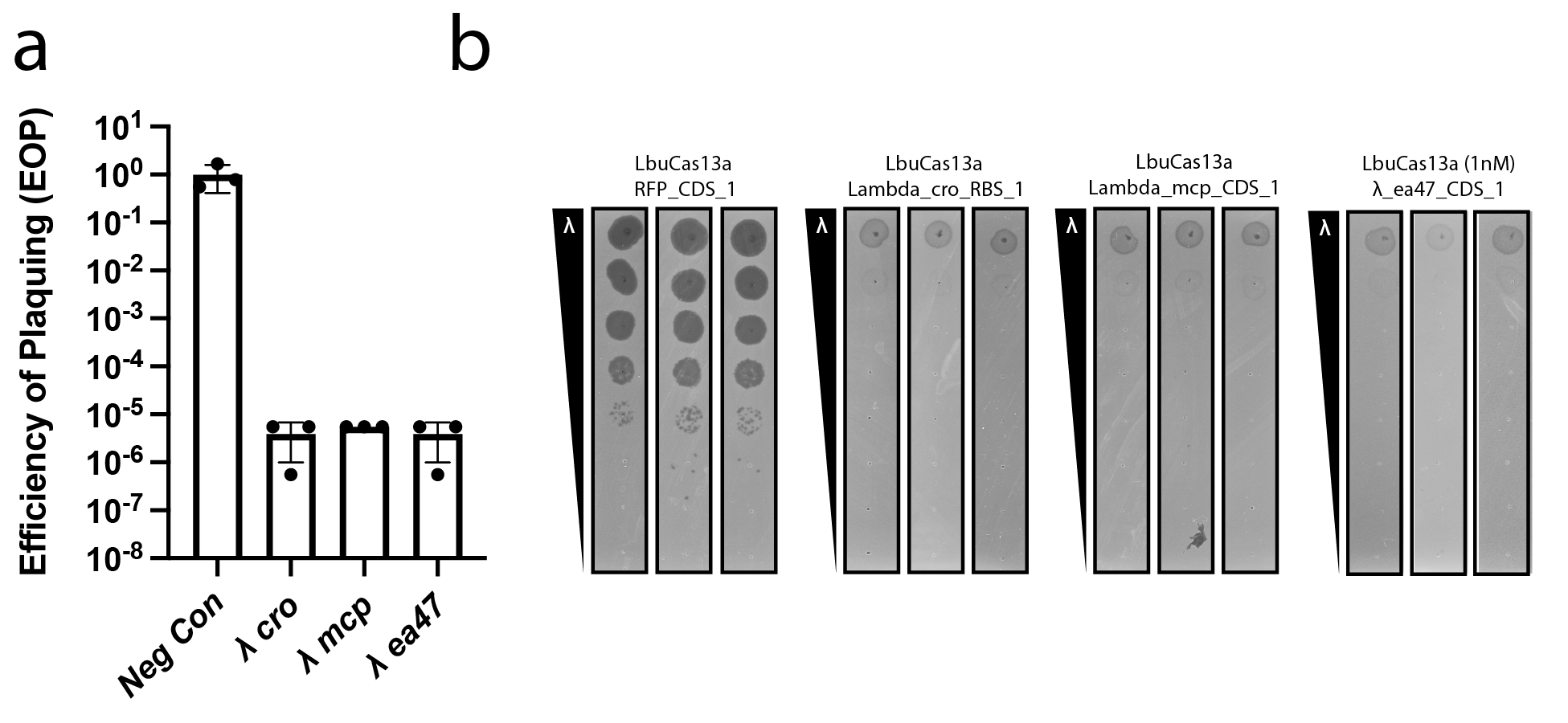

**Supplementary Fig. 5. Summary of results for LbuCas13a interference against phage λ.** **(a)** EOP estimation and **(b)** plaque assay results for λ infection against all crRNAs tested at 5 nM aTc induction of LbuCas13a. Due to toxicity at 5 nM aTc induction with LbuCas13a, crRNA Lambda_ea47_CDS_1 assays were performed at 1 nM aTc. For EOP graphs, results are represented as mean±std. All assays were performed in biological triplicate.

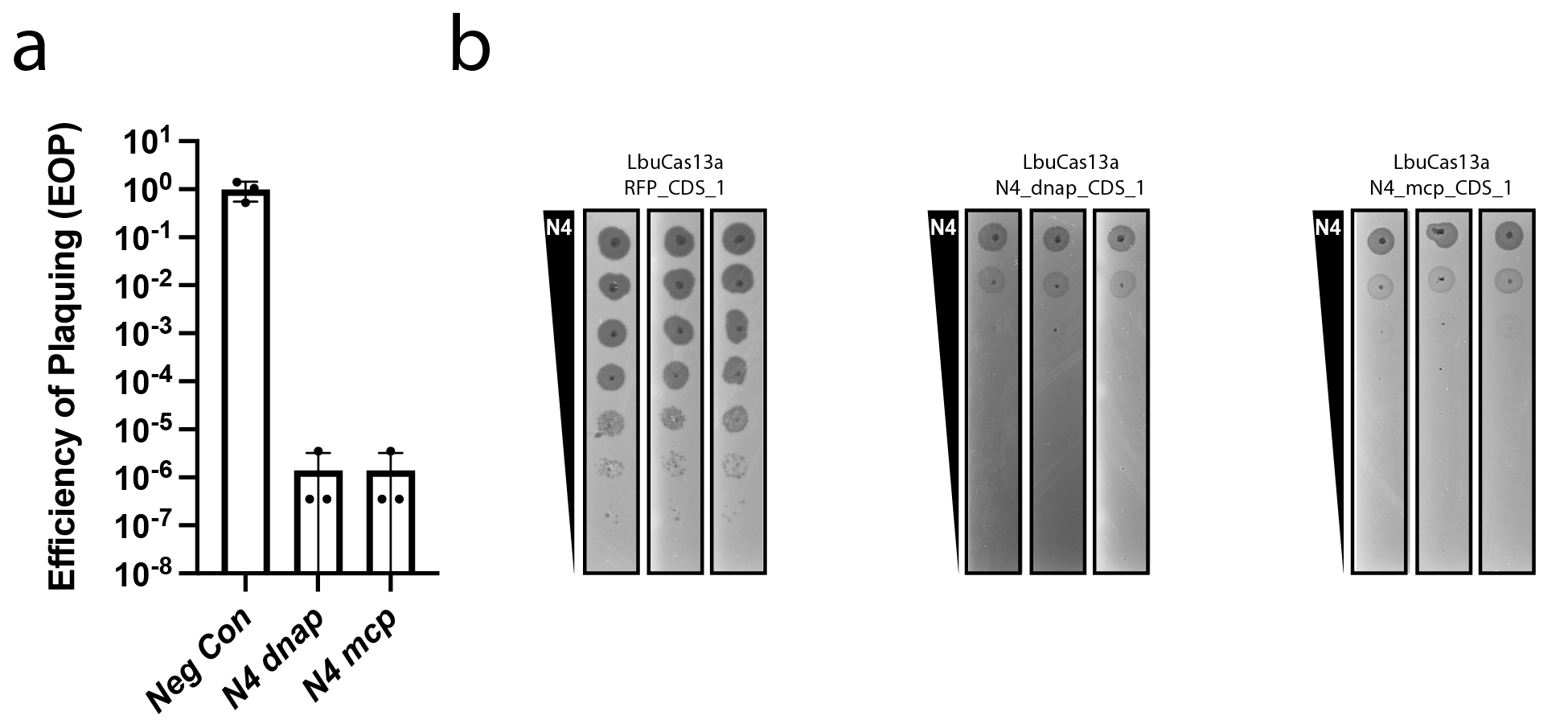

**Supplementary Fig. 6. Summary of results for LbuCas13a interference against phage N4.** **(a)** EOP estimation and **(b)** plaque assay results for N4 infection against all crRNAs tested at 5 nM aTc induction of LbuCas13a. For EOP graphs, results are represented as mean±std. All assays were performed in biological triplicate.

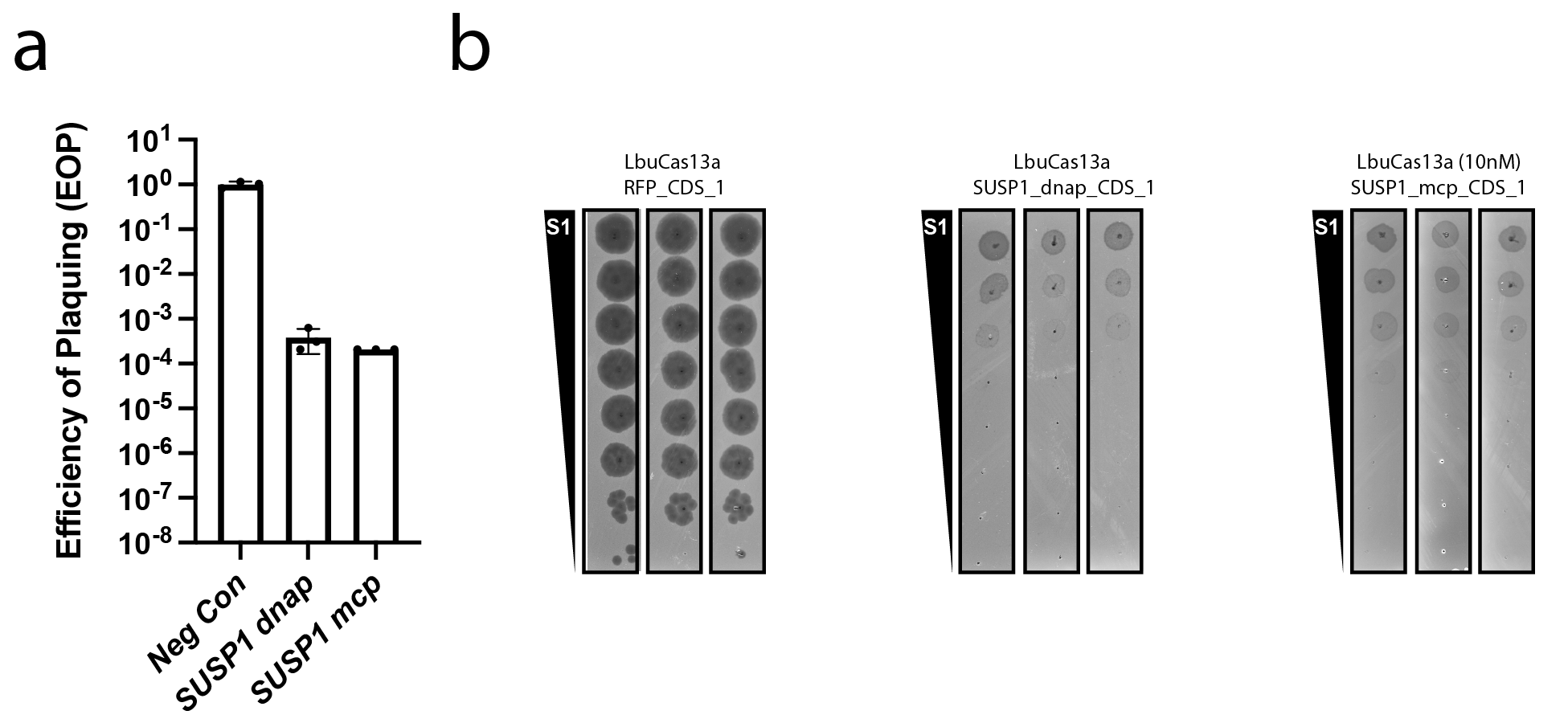

**Supplementary Fig. 7. Summary of results for LbuCas13a interference against phage SUSP1.** **(a)** EOP estimation and **(b)** plaque assay results for SUSP1 infection against all crRNAs tested at 5 nM aTc induction of LbuCas13a. For EOP graphs, results are represented as mean±std. All assays were performed in biological triplicate.

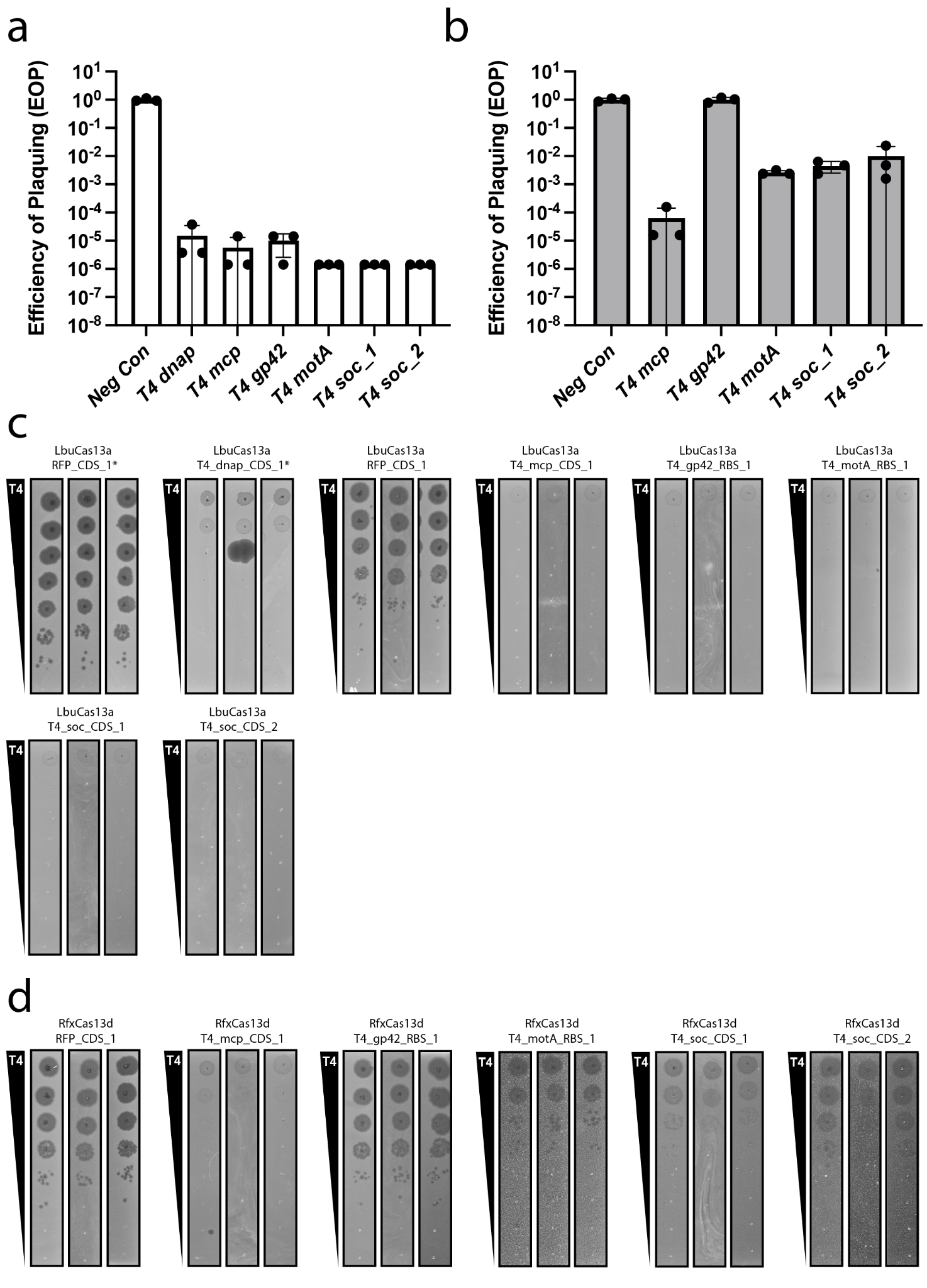

**Supplementary Fig. 8. Summary of results for LbuCas13a and RfxCas13d interference against phage T4.** **(a)** EOP estimation for T4 infection against all crRNAs tested at 5 nM aTc induction of LbuCas13a. **(b)** EOP estimation for T4 infection against all crRNAs tested at 5 nM aTc induction of RfxCas13d. **(c)** Plaque assay results for T4 infection against all crRNAs tested at 5 nM aTc induction of LbuCas13a. Experiments denoted with a * refer to a new preparation of T4 lysate. **(d)** Plaque assay results for T4 infection against all crRNAs tested at 5 nM aTc induction of RfxCas13d. For EOP graphs, results are represented as mean±std. All assays were performed in biological triplicate.

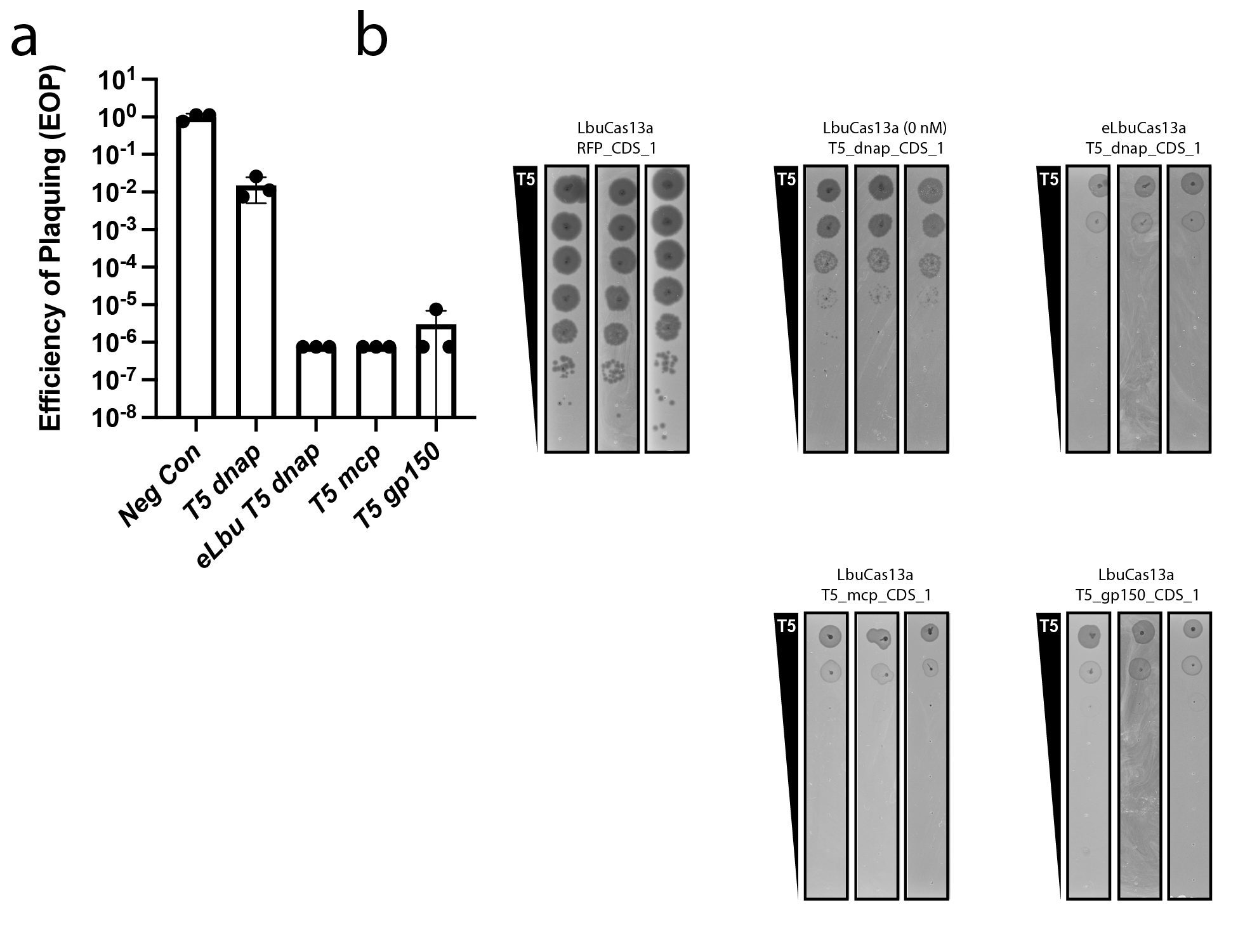

**Supplementary Fig. 9. Summary of results for LbuCas13a interference against phage T5.** **(a)** EOP estimation and **(b)** plaque assay results for T5 infection against all crRNAs tested at 5 nM aTc induction of LbuCas13a unless stated otherwise. Due to toxicity at 5 nM aTc induction with LbuCas13a, crRNA T5_dnap_CDS_1 assays were performed at 0 nM aTc. Additionally crRNA_dnap_CDS_1 was cloned into a reduced toxicity variant of LbuCas13a referred to as eLbuCas13a (Methods), which is also shown at 5 nM aTc induction. For EOP graphs, results are represented as mean±std. All assays were performed in biological triplicate.

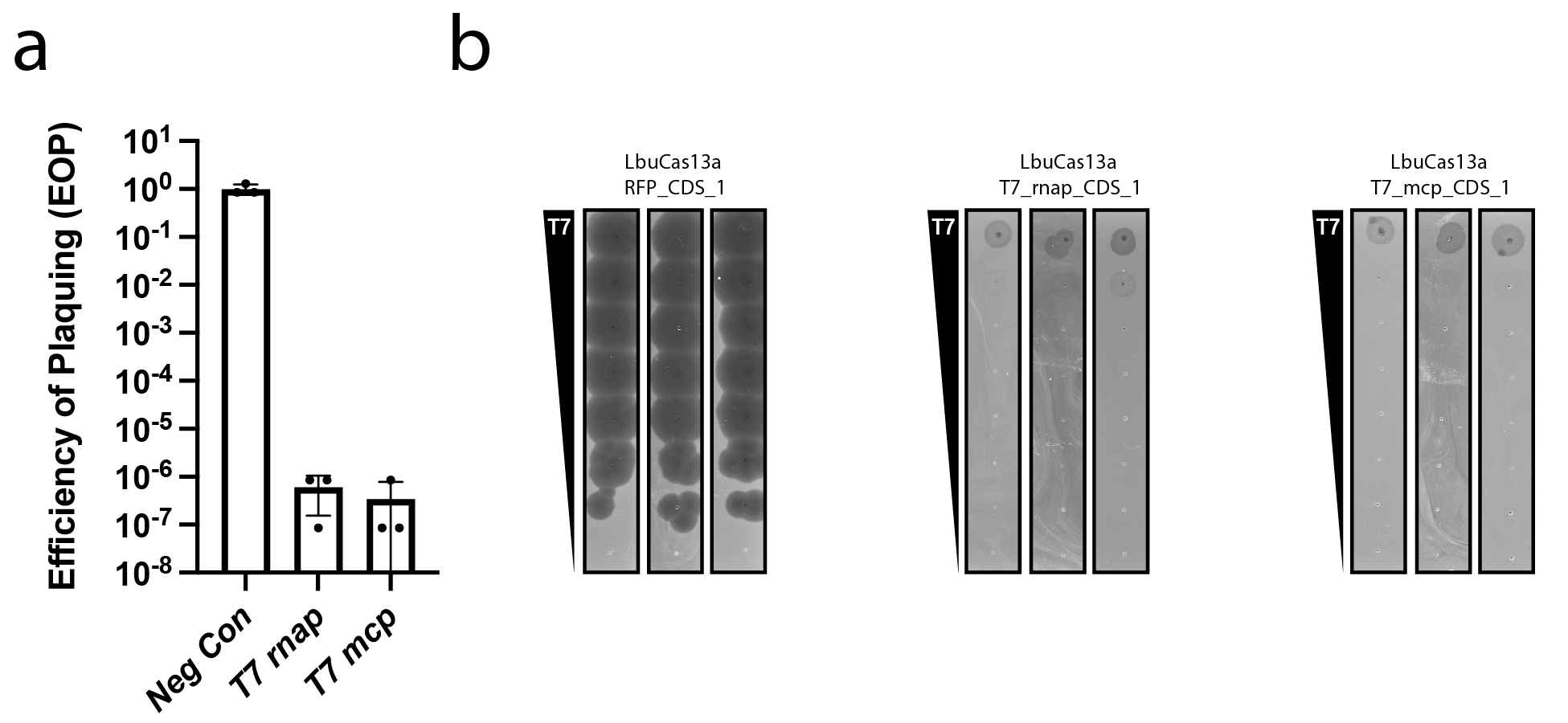

**Supplementary Fig. 10. Summary of results for LbuCas13a interference against phage T7.** **(a)** EOP estimation and **(b)** plaque assay results for T7 infection against all crRNAs tested at 5 nM aTc induction of LbuCas13a. For EOP graphs, results are represented as mean±std. All assays were performed in biological triplicate.

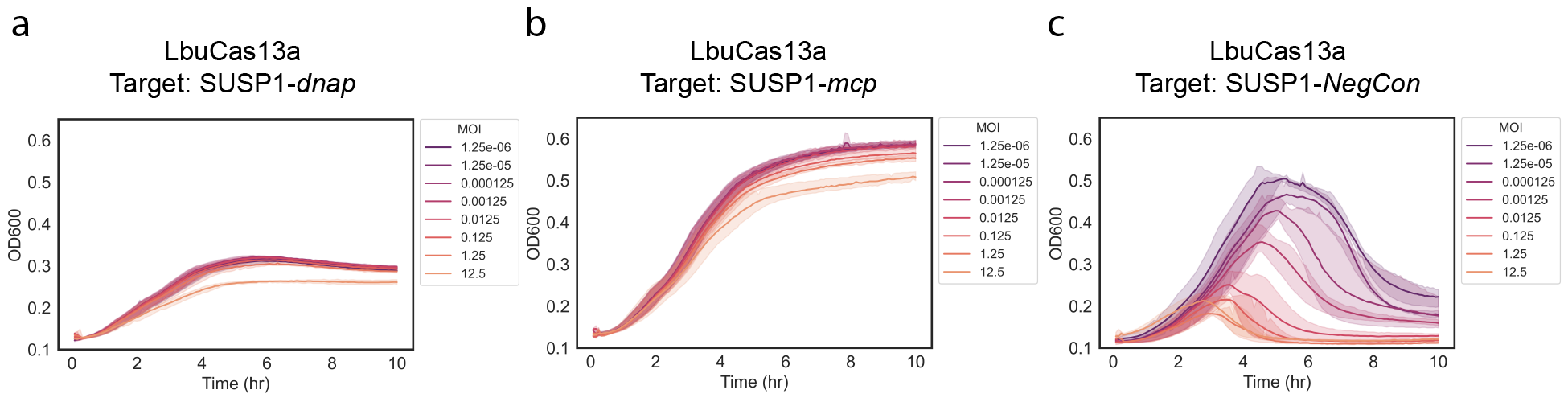

**Supplementary Fig. 11. SUSP1 displays sensitivity to LbuCas13a targeting during liquid infection.** Liquid infection experiments for phage SUSP1 against *E.coli* expressing LbuCas13a and a crRNA targeting **(a)** SUSP1*dnap*, **(b)** SUSP1*mcp*, and **(c)** a non-targeting crRNA across a range of MOIs. All assays were performed at 10 nM aTc induction with 3 biological replicates.

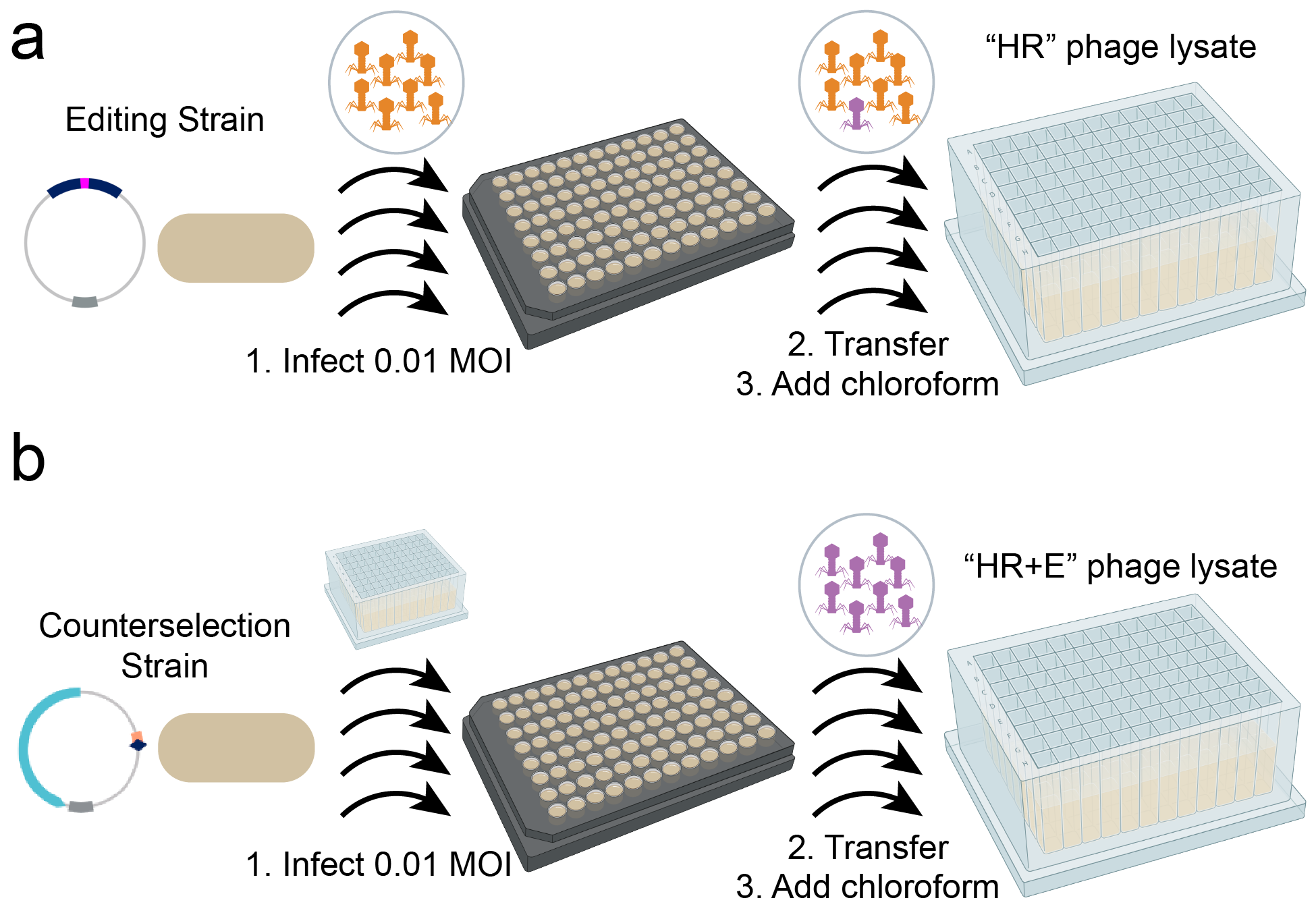

**Supplementary Fig. 12. Method overview for Cas13a Phage Editing. (a)** Homologous Recombination edited (HR) phage lysate is created by infection of an editing strain at 0.01 MOI. After lysis is observed phage lysates comprising a mixture of edited (purple) and wildtype (orange) phage are transferred to a deep well block and a drop of chloroform is added. These are the “HR” phage lysates. **(b)** Enriched phage lysates are created by infection of an counterselection strain at 0.01 MOI from an “HR” phage lysate. After lysis is observed phage lysates comprising primarily of edited (purple) phage are transferred to a deep well block and a drop of chloroform is added. These are the “HR+E” phage lysates.

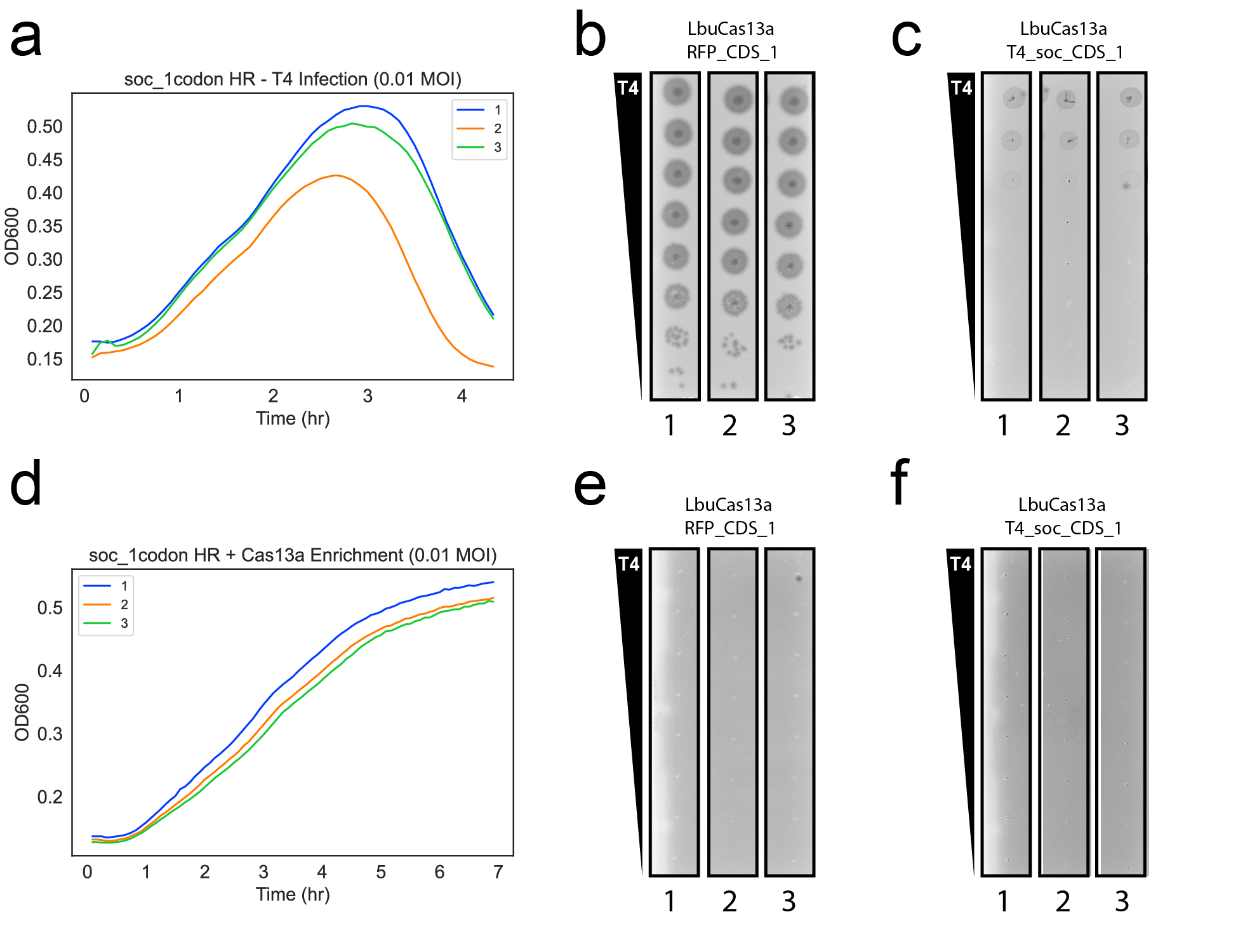

**Supplementary Fig. 13. Overview of phenotypic results for editing attempt *soc*-C. (a)** Growth curves for the homologous recombination editing step for *soc*-C. **(b)** Non-selective plaquing assay for lysates from (a) on a non-targeting crRNA. **(c)** Selective plaquing assay for lysates from (a) using the corresponding *soc*-targeting crRNA. **(d)** Growth curves for the enrichment step for lysates harboring, but not enriched for the *soc*-C edit. **(e)** Non-selective plaquing assay for lysates from (d) on a non-targeting crRNA. **(f)** Selective plaquing assay for lysates from (d) using the corresponding *soc*-targeting crRNA. For all liquid culture and plate-based assays shown using Cas13a, induction was performed with 10 nM aTc. Labels 1-3 correspond to unique parallel editing workflows.

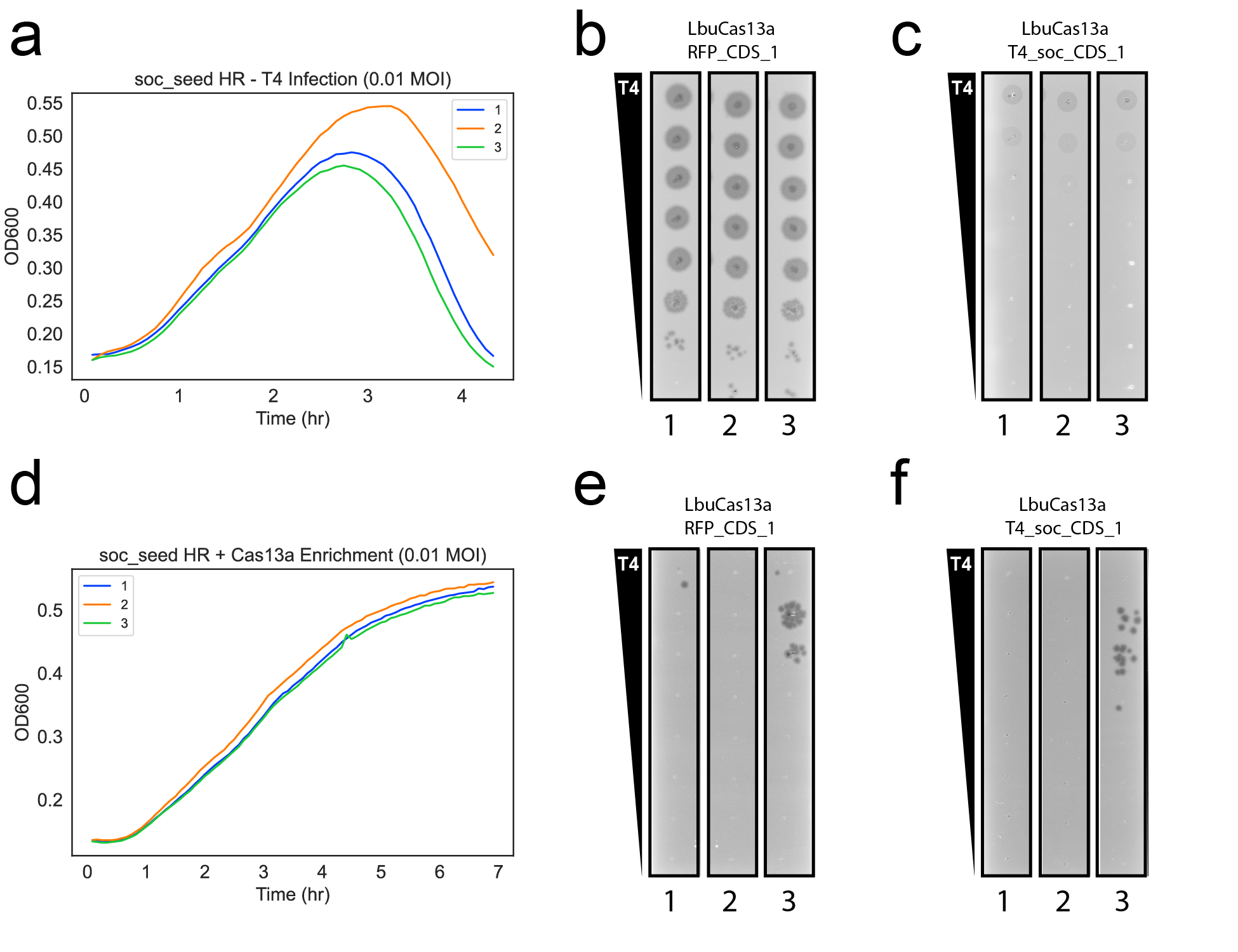

**Supplementary Fig. 14. Overview of phenotypic results for editing attempt *soc*-S. (a)** Growth curves for the homologous recombination editing step for *soc*-S. **(b)** Non-selective plaquing assay for lysates from (a) on a non-targeting crRNA. **(c)** Selective plaquing assay for lysates from (a) using the corresponding *soc*-targeting crRNA. **(d)** Growth curves for the enrichment step for lysates harboring, but not enriched for the *soc*-S edit. **(e)** Non-selective plaquing assay for lysates from (d) on a non-targeting crRNA. **(f)** Selective plaquing assay for lysates from (d) using the corresponding *soc*-targeting crRNA. For all liquid culture and plate-based assays shown using Cas13a, induction was performed with 10 nM aTc. Labels 1-3 correspond to unique parallel editing workflows.

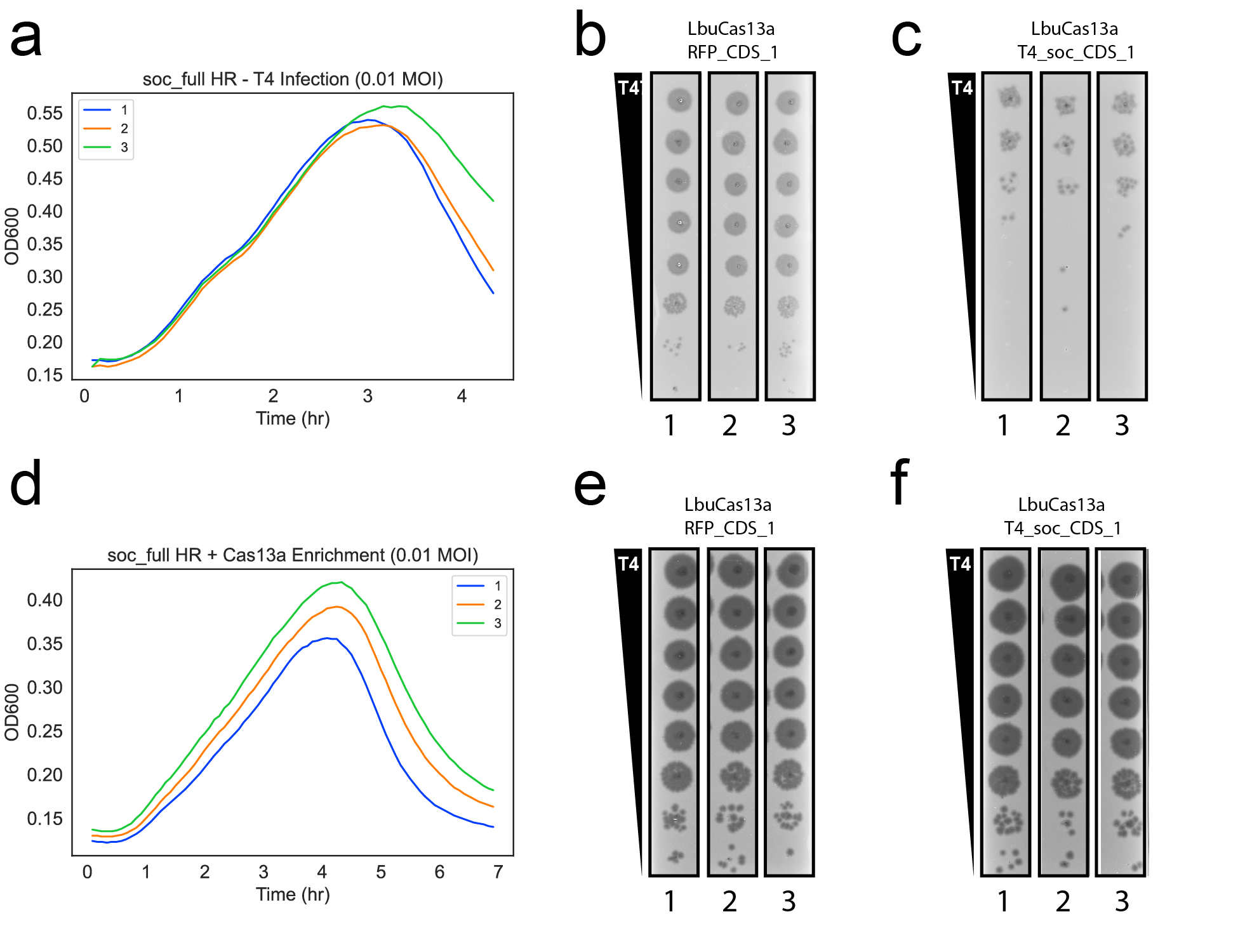

**Supplementary Fig. 15. Overview of phenotypic results for editing attempt *soc*-F. (a)** Growth curves for the homologous recombination editing step for *soc*-F. **(b)** Non-selective plaquing assay for lysates from (a) on a non-targeting crRNA. **(c)** Selective plaquing assay for lysates from (a) using the corresponding *soc*-targeting crRNA. **(d)** Growth curves for the enrichment step for lysates harboring, but not enriched for the *soc*-F edit. **(e)** Non-selective plaquing assay for lysates from (d) on a non-targeting crRNA. **(f)** Selective plaquing assay for lysates from (d) using the corresponding *soc*-targeting crRNA. For all liquid culture and plate-based assays shown using Cas13a, induction was performed with 10 nM aTc. Labels 1-3 correspond to unique parallel editing workflows.

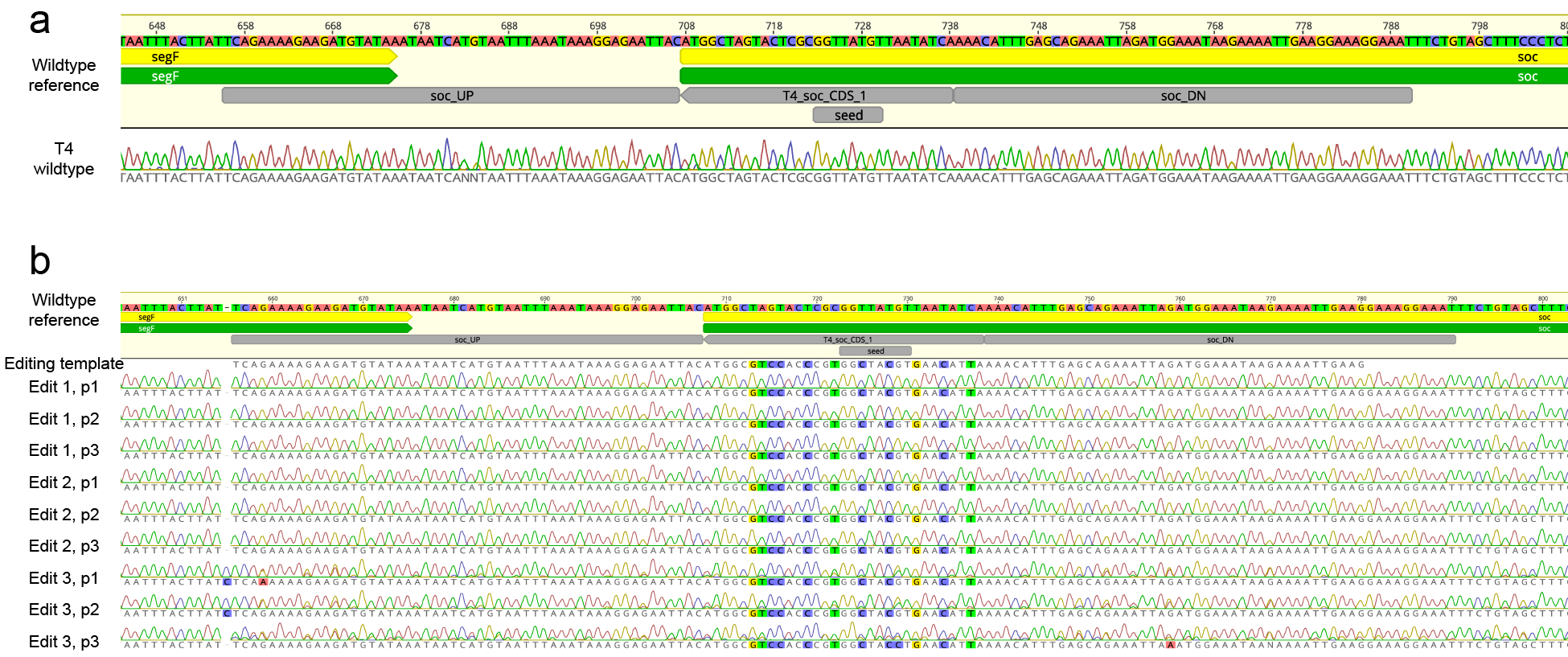

**Supplementary Fig. 16. Overview of genotyping results for *soc* editing attempts. (a)** Sanger sequencing trace from a PCR at the wildtype T4 *soc* locus. **(b)** Sanger sequencing trace from unbiased PCRs on individual plaques at the *soc* locus for *soc-F* edits. Results for three individual plaques (p1, p2, p3) are shown for enriched, selective plaquing from three independent editing processes (for instance 3 plaques from each of the dilutions shown in Supplementary Fig. 15f). The wildtype T4*soc* locus and the editing template is shown as a reference. Mutations relative to the wildtype locus are highlighted.

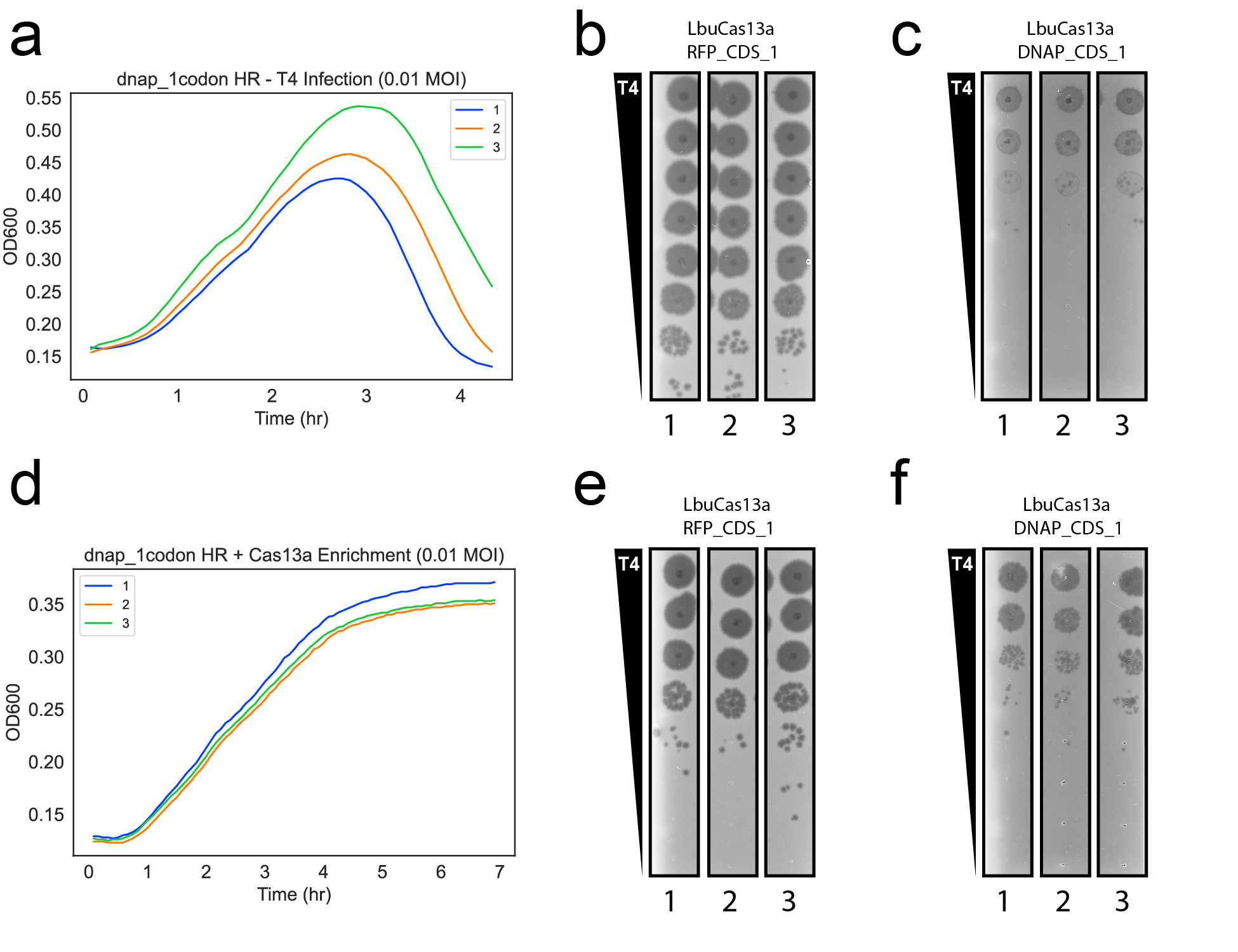

**Supplementary Fig. 17. Overview of phenotypic results for editing attempt *dnap*-C. (a)** Growth curves for the homologous recombination editing step for *dnap*-C. **(b)** Non-selective plaquing assay for lysates from (a) on a non-targeting crRNA. **(c)** Selective plaquing assay for lysates from (a) using the corresponding *dnap*-targeting crRNA. **(d)** Growth curves for the enrichment step for lysates harboring, but not enriched for the *dnap*-C edit. **(e)** Non-selective plaquing assay for lysates from (d) on a non-targeting crRNA. **(f)** Selective plaquing assay for lysates from (d) using the corresponding *dnap*-targeting crRNA. For all liquid culture assays shown using Cas13a, induction was performed with 10 nM aTc. For all plaquing assays shown using Cas13a, induction was performed with 5 nM aTc. Labels 1-3 correspond to unique parallel editing workflows.

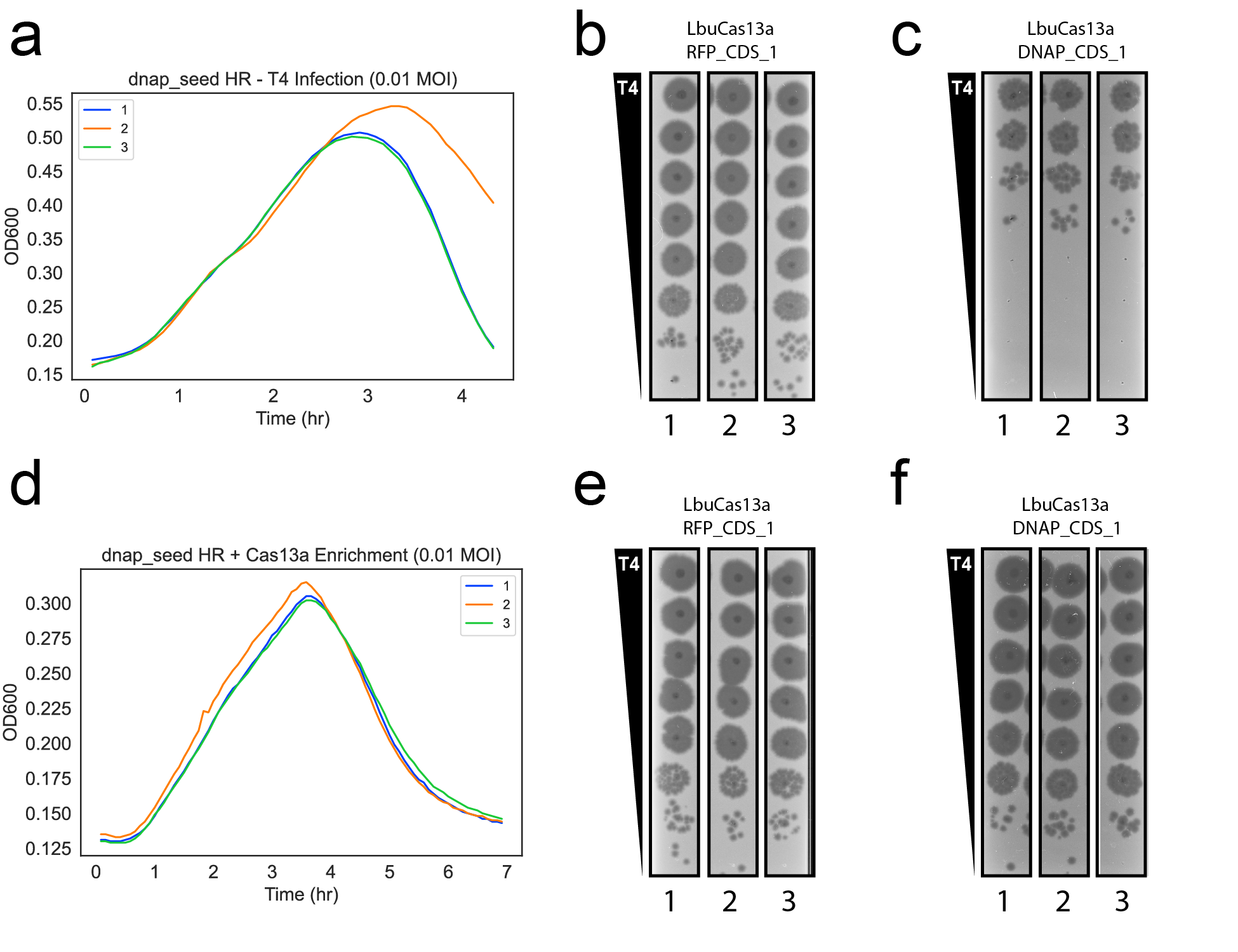

**Supplementary Fig. 18. Overview of phenotypic results for editing attempt *dnap*-S. (a)** Growth curves for the homologous recombination editing step for *dnap*-S. **(b)** Non-selective plaquing assay for lysates from (a) on a non-targeting crRNA. **(c)** Selective plaquing assay for lysates from (a) using the corresponding *dnap*-targeting crRNA. **(d)** Growth curves for the enrichment step for lysates harboring, but not enriched for the *dnap*-S edit. **(e)** Non-selective plaquing assay for lysates from (d) on a non-targeting crRNA. **(f)** Selective plaquing assay for lysates from (d) using the corresponding *dnap*-targeting crRNA. For all liquid culture assays shown using Cas13a, induction was performed with 10 nM aTc. For all plaquing assays shown using Cas13a, induction was performed with 5 nM aTc. Labels 1-3 correspond to unique parallel editing workflows.

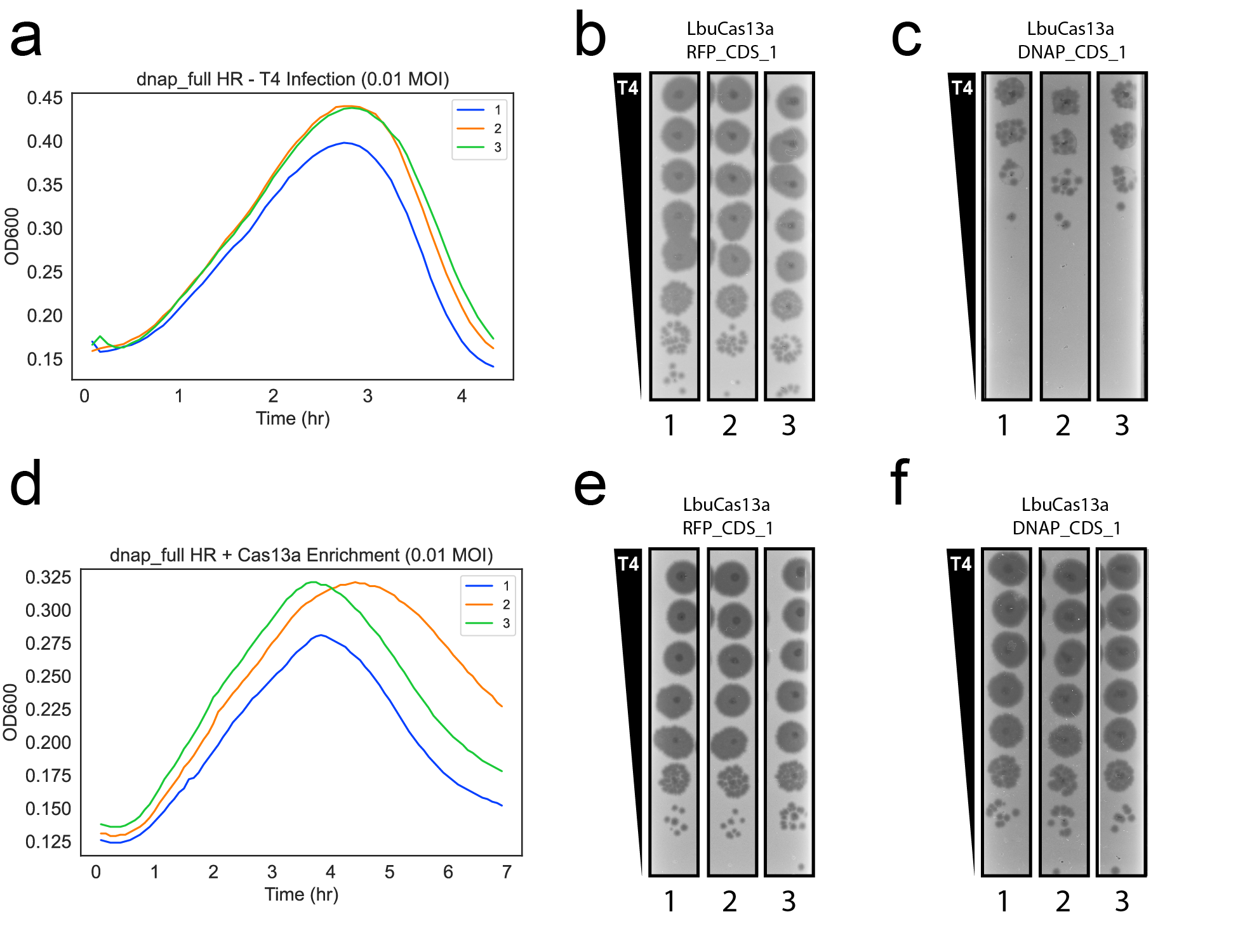

**Supplementary Fig. 19. Overview of phenotypic results for editing attempt *dnap*-F. (a)** Growth curves for the homologous recombination editing step for *dnap*-F. **(b)** Non-selective plaquing assay for lysates from (a) on a non-targeting crRNA. **(c)** Selective plaquing assay for lysates from (a) using the corresponding *dnap*-targeting crRNA. **(d)** Growth curves for the enrichment step for lysates harboring, but not enriched for the *dnap*-F edit. **(e)** Non-selective plaquing assay for lysates from (d) on a non-targeting crRNA. **(f)** Selective plaquing assay for lysates from (d) using the corresponding *dnap*-targeting crRNA. For all liquid culture assays shown using Cas13a, induction was performed with 10 nM aTc. For all plaquing assays shown using Cas13a, induction was performed with 5 nM aTc. Labels 1-3 correspond to unique parallel editing workflows.

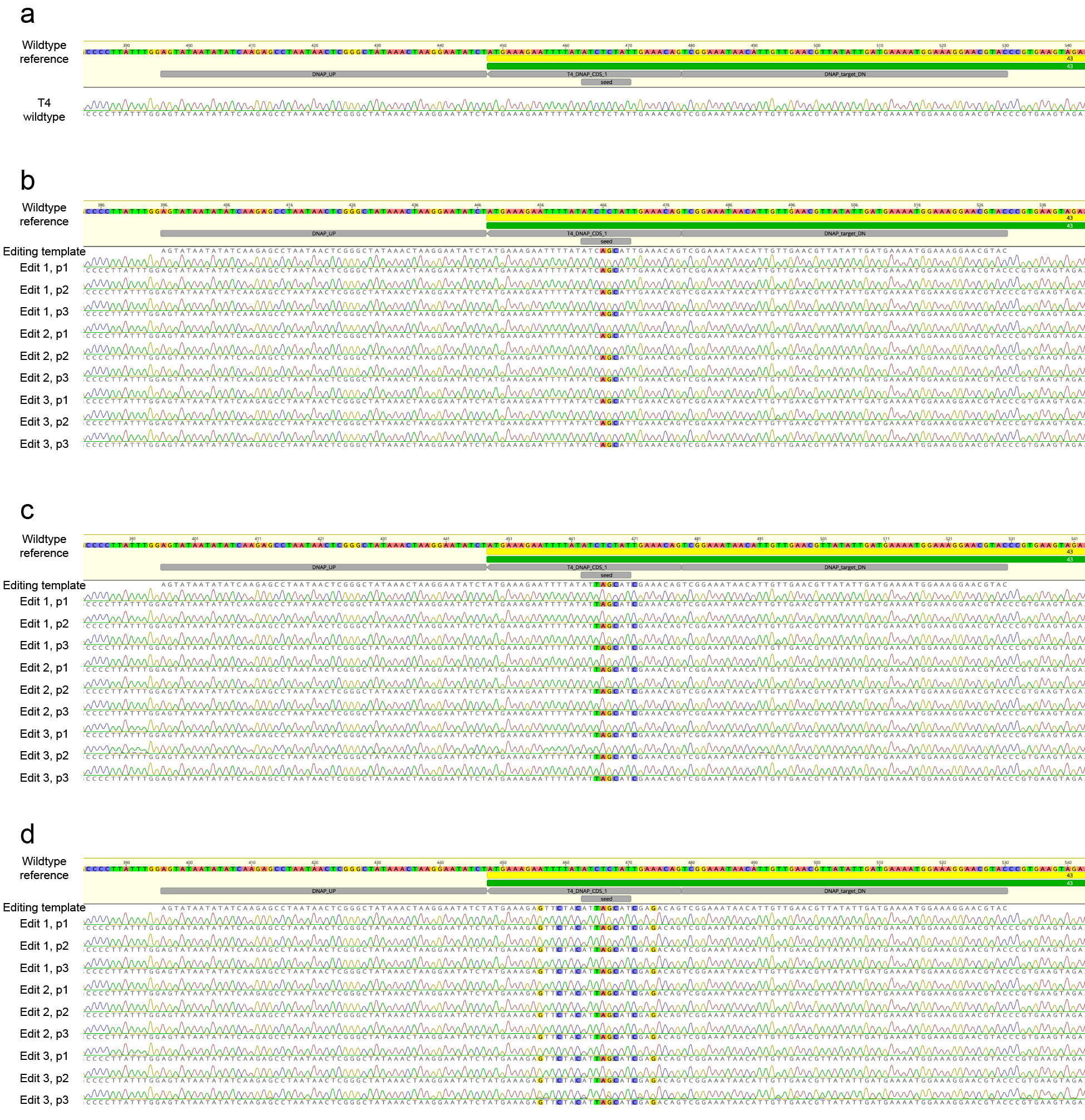

**Supplementary Fig. 20. Overview of genotyping results for *dnap* editing attempts. (a)** Sanger sequencing trace from a PCR at the wildtype T4 *dnap* locus. **(b-d)** Sanger sequencing trace from unbiased PCRs on individual plaques at the *dnap* locus for **(b)** *dnap*-C, **(c)** *dnap*-S, and **(d)** *dnap*-F edits. Results for three individual plaques (p1, p2, p3) are shown for enriched, selective plaquing from three independent editing processes (for instance 3 plaques from each of the dilutions shown in Supplementary Fig. 17f, 18f, and 20f for *dnap*-C, *dnap*-S, and *dnap*-F respectively). The wildtype T4*dnap* locus and the editing template is shown as a reference. Mutations relative to the wildtype locus are highlighted.
